# Vertical profile of airborne microbial communities in the Southern Ocean atmospheric boundary layer

**DOI:** 10.64898/2026.08.26.747214

**Authors:** Sofía Galbán, Woo Young Kim, Pablo Sanz, Tamara Pletzer, Manuel Bañón, Juan Antonio Higuera, Javier Méndez, Kang-ho Ahn, Sergi González-Herrero, Ana Justel, Antonio Quesada

**Author notes:** These authors contributed equally to this work (Sergi González-Herrero, Ana Justel, Antonio Quesada).

## Abstract

Aerobiological studies have largely focused on near-surface sampling and horizontal biogeographic patterns, while vertical structuring of airborne microbial communities within the atmospheric boundary layer (ABL) remains poorly understood. Here, we investigated microbial communities across the lower and upper ABL in a low-orography coastal site on the Antarctic Peninsula, representative of the Southern Ocean marine ABL and with low direct human influence. Airborne microorganisms were sampled simultaneously using ground-based and aerial platforms on five occasions. Community composition, abundance, and cell morphometry were assessed using metabarcoding and epifluorescence microscopy and interpreted alongside atmospheric observations. Airborne bacterial and eukaryotic communities showed consistent vertical stratification, although partial taxonomic overlap indicates vertical connectivity between atmospheric layers. Lower ABL communities were more diverse than upper ABL counterpart, compositionally homogeneous, and dominated by marine-associated taxa, reflecting strong influence from local sources and turbulent mixing. In contrast, upper ABL communities were less diverse but more heterogeneous among sampling events, enriched in stress-tolerant, terrestrial and plant-associated taxa, consistent with atmospheric filtering, selective upward transport, and long-range atmospheric inputs. Upper-layer samples also exhibited higher microbial abundance and greater prevalence of elongated cell morphologies, suggesting particle accumulation aloft and aerodynamic selection permanence. Together, these findings identify the Southern Ocean ABL as a vertically structured microbial habitat organized into two partially decoupled sublayers, in which atmospheric dynamics regulate microbial dispersal, ecosystem connectivity, and biogeographic patterns.

## INTRODUCTION

The atmosphere is increasingly recognized as a dynamic microbial environment that connects land and sea habitats at multiple spatial scales through the aerial dispersal of microbial cells, propagules, and biomolecules. Despite growing interest in the aeromicrobiome over the past decade, our understanding remains comparatively limited in both scope and depth relative to other ecosystems [1].

This environment is mainly structured into vertical layers, whose altitude and thickness are primarily determined by thermal properties [2]. The troposphere, extending from the Earth’s surface up to approximately 10-20 km, contains the bulk of atmospheric mass, including most of the bioaerosols [3]. The atmospheric boundary layer forms its lowest sublayer, spanning from the surface to ∼0.3–3 km above the ground. Its upper limit is defined by a thin temperature inversion layer marking the transition to the free troposphere, which is characterised by more stable, stratified, and laminar flow [4]. Within the ABL, turbulent mixing driven by wind shear and surface heating promotes strong vertical exchange of particles and bioaerosols between the surface and the atmosphere [3].

Most aerobiology studies have focused primarily on horizontal spatial patterns and near-ground temporal dynamics of airborne microbial communities, while vertical variations across different atmospheric heights remain poorly understood. In particular, microbial data from the upper ABL and the free troposphere are scarce due to logistical and methodological challenges, including limited accessibility, reliance on aerial sampling platforms (such as towers, unmanned vehicles, tethered balloons or aircrafts), constraints on sufficient air sampling volume, strict contamination control, and highly variable meteorological conditions [1,5]. The few vertical aerobiological studies conducted to date have largely focused on urban environments [6,7], dust-associated transport events [8], and ecosystem-specific processes in forested environments [9,10]. However, most of these studies do not consistently employ simultaneous multi-height sampling, limiting their ability to understand short-term vertical variability.

The polar regions are environments of high ecological and biogeographical relevance for studying vertical bioaerosol dynamics, due to minimal local anthropogenic influence (i.e., absence of human associated microbes and far from human driven aerosolization processes) enables the observation of bioaerosols under conditions of reduced human impact [11,12]. Antarctica is uniquely isolated from other land masses, with long-range atmospheric transport representing the direct pathway by which airborne microorganisms could reach the continent. The persistent katabatic wind regime surrounding the continent strongly limits low-level air exchange with the surrounding Southern Ocean. The main entry pathway for externally sourced microorganisms is through the air at hundred meters aloft where air masses typically reach the continent via secondary circulations [13]. Despite the potential importance of the upper atmosphere as a transport pathway, vertical microbial profiles in polar regions remain extremely poorly resolved, as sampling of airborne microorganisms is further challenged by low microbial biomass and severe logistical constraints, resulting in a pronounced data gap. To date, and, as far as we know, only two publications [14,15] have characterised vertical profiles of airborne microbial communities in Antarctica using simultaneous upper and lower ABL sampling at Syowa station (coastal area) and S17 base (continental ice sheet). Both studies represent a single-event observation and therefore cannot elucidate the importance of the atmospheric condition variability in airborne bioaerosols. Archer et al. (2019) [16] also examined Antarctic airborne microbial communities using near-surface and ∼2000 m air samples, but these were not collected simultaneously and the study focused on microbial isolation and atmospheric dispersal rather than ABL vertical gradients. By contrast, vertical studies of non-biological aerosols in polar regions, although still scarce, are more numerous, providing a better-resolved physical framework of their vertical structure [17,18]. Overall, the scarcity of simultaneous multi-height measurements of airborne microbial communities in polar regions considerably restricts the ability to resolve the role of long-range atmospheric dispersal in microbial colonization and therefore the potential establishment of communities in newly deglaciated environments.

In this study, we explored the temporal and vertical variability of the microorganisms in Byers Peninsula located in Livingston Island, an Antarctic island in the Southern Ocean. The location is far from upstream land masses and from any direct human influence, which allows to understand the mechanisms underlying the aerobiological short and long-range transport. Using a sampling tower and tethered balloon simultaneously for several days, combined with metabarcoding and fluorescence microscopy, and supported by atmospheric observations and air mass back-trajectory analyses, we showed that all lines of evidence consistently converge on a vertical structuring of airborne microorganisms across taxonomic, ecological, and morphological features. This pattern was reflected in a decline in diversity and strong differentiation between lower and upper ABL layers at different taxonomic ranks. Near-surface communities were more homogeneous and dominated by local marine inputs, whereas upper-layer communities were more variable and enriched in stress-tolerant terrestrial taxa, consistent with selective upward transport and greater influence of long-range atmospheric transport. This vertical organization was also evidenced by morphological features, with upper layer showing higher concentrations and a greater contribution of elongated cells. Overall, these patterns revealed that the ABL over Antarctic coastal regions constitutes a vertically organized system. They also support a conceptual framework in which a lower layer, dominated by turbulent mixing and local sources, transitions into an upper layer shaped by the history of transport and selective filtering acting on microbial traits that determine persistence and dispersion in the atmosphere.

## MATERIAL AND METHODS

### Sampling site, protocol and instrumentation

The study was conducted at Byers Peninsula, located at the western end of Livingston Island in the South Shetland Islands, Antarctic Peninsula (Fig. S1). This region was selected for its proximity to the sea, with low elevation, gentle topography, horizontal relief homogeneity and limited surface roughness [19], which together minimize local effects that might complicate the interpretation of the ABL structure [20]. At the same time, its small landmass (84.7 km^2^) and coastal setting favour strong interactions between marine and terrestrial air masses under atmospheric conditions representative of the Southern Ocean, where its summertime marine atmospheric boundary layer (MABL) is typically stable and vertically decoupled [21]. In addition, Byers Peninsula is designated as an Antarctic Specially Protected Area (ASPA No. 126) and therefore is an ideal setting for aerobiology studies because of the minimal direct human impact [22].

Sampling of airborne microbial communities was carried out on five different days during January 2023 using a ground-based sampling tower and a tethered balloon deployed simultaneously on the southern beach of the peninsula, approximately 0.4 km from Byers Camp (Fig. S1; Dataset S1). The site was inhabited by six people and no vehicles or vessels were present in the vicinity during sampling, minimizing potential contamination derived from human actions. Full technical description of the tethered balloon and the sampling tower setups together with the collecting protocol are provided in Supplementary Methods. Balloon- and tower-derived samples are labelled “B_” and “T_”, respectively.

### DNA extraction, amplification and sequence processing

Microbial cells were recovered from MicroAirCollectors and genomic DNA was extracted using the DNAeasy PowerSoil Kit (QIAGEN)® following Parro et al. (2025) [23]. Bacterial communities were characterized by amplification of the V3-V4 regions of the 16S rRNA gene using primers 341F/805R [24], whereas eukaryotic communities were assessed through amplification of the V9 region of the 18S rRNA gene using EMP primers 1391F/EukBr [25, 26]. Libraries were sequenced on an Illumina MiSeq platform (Microomics, Spain).

Sequence data were processed with DADA2 [27] to infer amplicon sequence variants (ASVs). Taxonomic assignment was performed against the SILVA v138.2 database [28] using naïve Bayesian classifiers implemented in DADA2 [29] and QIIME2 [30] for bacterial and eukaryotic datasets, respectively. Non-target sequences and ASVs detected in negative controls were removed prior to downstream analyses following conservative method for low biomass samples [31]. All procedures were performed under sterile conditions following best practises for low-biomass samples [31]. Detailed descriptions of contamination control procedures, sequence filtering, taxonomic curation and dataset processing are provided in the Supplementary Methods.

### Epifluorescence microscopy analysis

Airborne microbial cells extracted from the collectors, following Parro et al. (2025) [23], were stained with SYBR Green I and filtered onto 0.2 μm black polycarbonate membranes for epifluorescence microscopy analysis. Microbial abundance and morphometric characterization were determined using a Nikon Eclipse 80i microscope equipped with FITC fluorescence filters optimized for SYBR Green I detection. Cell counts were obtained from randomly selected fields of view and microorganisms were classified into size and morphotype categories following Galbán et al. (2021) [32]. Cell concentrations were expressed as cells mO³ of air and the relative proportions of size and morphotype categories were expressed as percentages of the total counted cells. Negative controls (CNN and CNS) consistently showed no detectable fluorescence, confirming the absence of contamination.

Differences in microbial abundance between atmospheric layers were evaluated using two-way ANOVA, whereas differences in the relative contribution of size and morphotype categories were assessed using Wilcoxon signed-rank tests. Statistical significance was established at *p* < 0.05. Detailed microscopy procedures, image analyses and counting criteria are described in the Supplementary Methods.

### Microbial community analyses

Microbial diversity patterns for both bacterial and eukaryotic communities were assessed using alpha and beta diversity metrics. Alpha diversity was estimated using richness and Shannon indices, and differences between ABL layers were tested using Welch’s t-tests. Beta diversity was assessed using Bray–Curtis dissimilarities and visualized through non-metric multidimensional scaling (NMDS). Differences in community composition between layers and sampling events were evaluated using permutational multivariate analysis of variance (PERMANOVA; 999 permutations). Rarefaction analyses confirmed adequate sequencing depth for both bacterial and eukaryotic datasets. All statistical analyses and visualizations were performed in R v4.2.3 [33] using phyloseq [34], vegan [35] and tidyverse suite [36] packages.

A subset of core ASVs and layer-representative taxa was used to explore community consistency and layer-specific patterns. Indicator taxa analyses [37] were performed to identify taxa associated with each atmospheric layer. Community overlap among sampling events and atmospheric layers was assessed by comparing shared and exclusive taxa across datasets. Detailed definitions of core taxa, layer-representative taxa, and overlap analyses are provided in the Supplementary Methods.

### Boundary layer depth

The height of the ABL was estimated using the bulk Richardson number, which can be calculated from the meteorological data measured during the ascent and descent phases of the flights (Dataset S2) as:

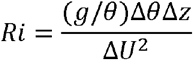

where g is the gravity constant in ms^−2^, z is the height in m, U is the horizontal wind speed in ms^−1^, and θ the potential temperature in K calculated as θ — T(1000/P)^0.286^, where T is the temperature in K and P is the air pressure in hPa. The increments (Δ) were calculated from the ground. The boundary layer height is considered the height where *Ri* > 0.25 [38].

The ABL height was not estimated for sampling event 1 due to the absence of concurrent balloon meteorological data required for *Ri* calculation. Samples from Balloons 2–4 represented air masses spanning the upper ABL, whereas sample from Balloon 5 represented consistently free tropospheric air (Fig. S2, S3). For simplicity, all samples from balloons are referred to as “upper ABL” throughout the manuscript. Further details on the definition of sampling heights are provided in the Supplementary Methods.

### Atmospheric transport analysis

Atmospheric transport was analysed using five-day air mass back-trajectories computed with the Hybrid Single-Particle Lagrangian Integrated Trajectory (HYSPLIT) model [39] based on ERA5 reanalysis data. To better constrain the atmospheric transport potential of airborne microorganisms, dry deposition was incorporated into the analyses following Galbán et al. (2021) [32], using the most representative particle sizes observed by microscopy for both upper- and lower-layer sampled air masses. In addition, Lagrangian coherent structures (LCS) were characterised using finite-time Lyapunov exponent (FTLE) analysis to identify large-scale attracting atmospheric transport features during sampling periods [40]. Detailed modelling settings and parameterisations are provided in the Supplementary Methods

## RESULTS

### Heterogeneity of the microbial vertical profile

This study detected a total of 9,484 bacterial ASVs (895,741 reads) and 989 eukaryotic ASVs (447,007 reads) across all sampling events, with a pronounced vertical imbalance in community richness and structure in both domains. For bacteria, 8,648 ASVs were identified in the lower ABL and 877 ASVs in the upper ABL, with significantly higher average richness in the lower layer (Fig. 1A). A significant similar pattern was observed for eukaryotes (893 vs. 137 ASVs; Fig. 1B). The average Shannon index was also significantly higher in the lower ABL for bacteria (Fig. 1C), whereas higher but no significant differences (*p* = 0.07) were observed for eukaryotes (Fig. 1D). Differences in community structure, both bacterial and eukaryotic domain, were supported by Bray-Curtis dissimilarities with NMDS ordination showing a clear and significant segregation between lower and upper ABL samples (Fig. 1E, F). Overlap between lower and upper ABL was minimal. Only 41 bacterial ASVs were shared, representing 12.1% of the total reads from the upper ABL bacterial community but just 1.93% of the lower. Similarly, 41 eukaryotic ASVs were shared, comprising a large fraction of the upper ABL community (82.2%) but a small proportion of the lower ABL (21.3%).

**Fig. 1.**
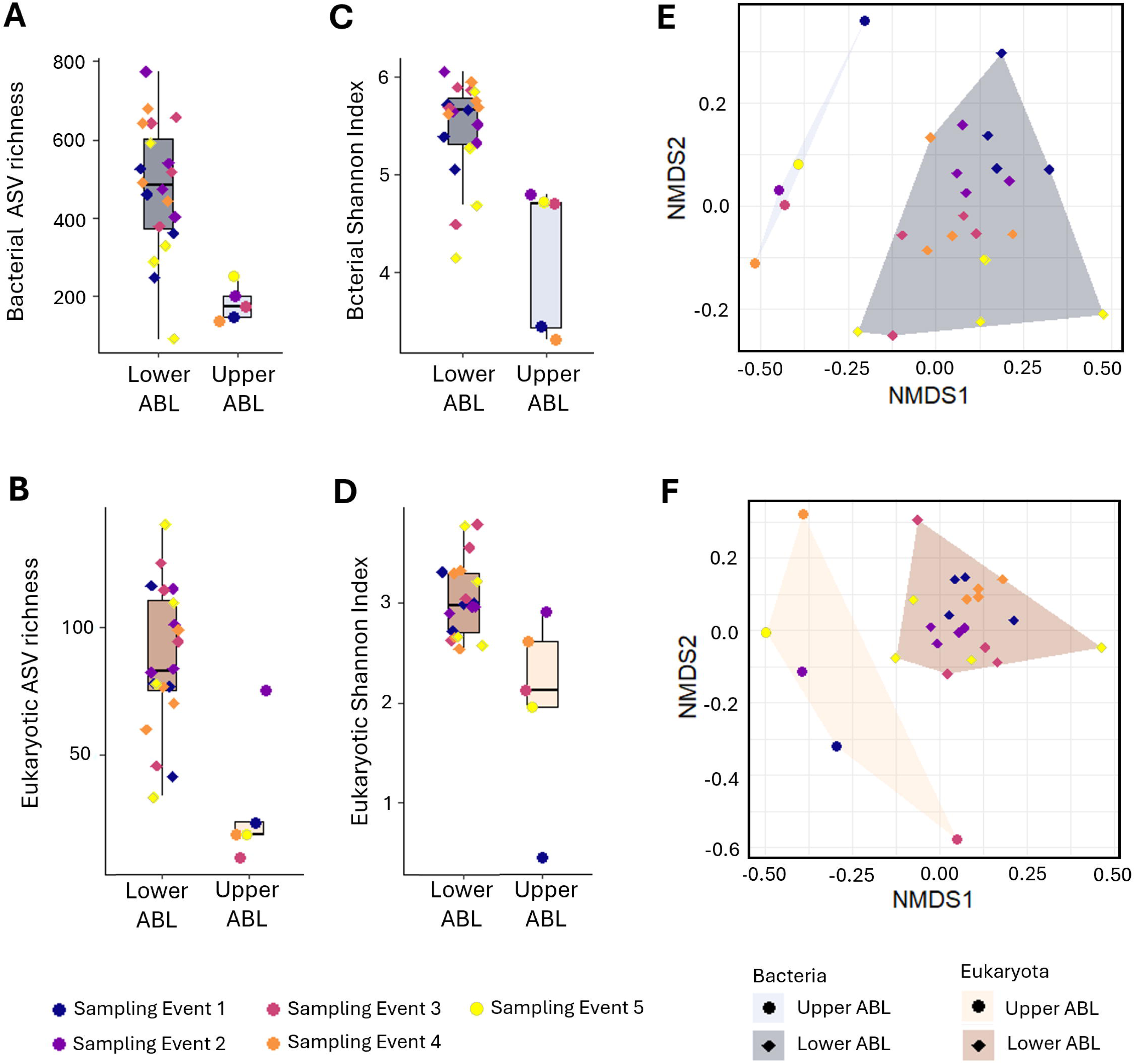
Airborne microbial community diversity and structure across the lower and upper ABL. **A-B.** Bacterial (**A**) and eukaryotic (**B**) ASV richness. Average richness for both domains differs significantly between the two ABL layers (Welch’s t-test, *p* < 0.05). **C-D.** Bacterial (**C**) and eukaryotic (**D**) Shannon Index. Bacterial Shannon Index differs significantly between ABL layers (Welch’s t-test, *p* < 0.05), whereas for eukaryote does not (Welch’s t-test, *p* = 0.07). **E-F.** NMDS ordination, based on Bray-Curtis dissimilarity of bacterial ASVs (**E**) and eukaryotic ASVs (**F**), showing the differences in community structure across ABL layers and sampling events. **E**. Bacterial community composition differs significantly between upper and lower ABL (PERMANOVA test, *p* ≤ 0.001), but not between sampling events (PERMANOVA test, *p* = 0.09). **F**. Eukaryotic community composition differs significantly between upper and lower ABL and between sampling events (PERMANOVA test, *p* ≤ 0.001).

At high altitude within the ABL, microbial communities showed pronounced heterogeneity across sampling events. Core community analysis identified only six bacterial and one eukaryotic ASVs shared by at least 60% of the samples (Core60), out of 877 and 137 ASVs detected in the upper ABL, respectively. For bacteria, the six bacterial Core60 ASVs included *Arthrobacter* and *Conexibacter*, genera associated with pristine soils (Dataset S3, S3), while the single eukaryotic ASV could not be confidently assigned at the phylum level (Dataset S3). These ASVs contributed unevenly across events. In contrast, lower ABL communities were highly homogeneous across events. A stable bacterial Core60 community (102 ASVs) contributed similarly to community relative abundance across samples (16.2-23.67%) and was largely composed of marine-associated genera (Dataset S5, S5). Eukaryotic communities in the lower ABL showed similar consistency, with 100 Core60 ASVs representing 76.49% of total eukaryotic reads, dominated by fungi and marine-associated algae and protist taxa (Dataset S5, S6).

To further explore the ecological organization of airborne bacterial communities across ABL layers, we examined distribution patterns at broader taxonomic levels. Across all samples, we detected 42 bacterial phyla, dominated by Actinomycetota, Pseudomonadota, Bacteroidota, Bacillota, Chloroflexota and Verrucomicrobiota. Within all bacterial phyla, 326 families and 153 uncultured families were detected across the vertical profile. Several bacterial families showed marked differences in abundance between layers (Fig. 2A), with 35 families enriched in the lower ABL and 32 in the upper ABL (Fig. S4A-B). Lower ABL communities were dominated by marine and Antarctic fauna-associated bacteria, whereas upper ABL communities were enriched in soil- and plant-associated taxa (Fig. 2B; Dataset S4, S5).

**Fig. 2.**
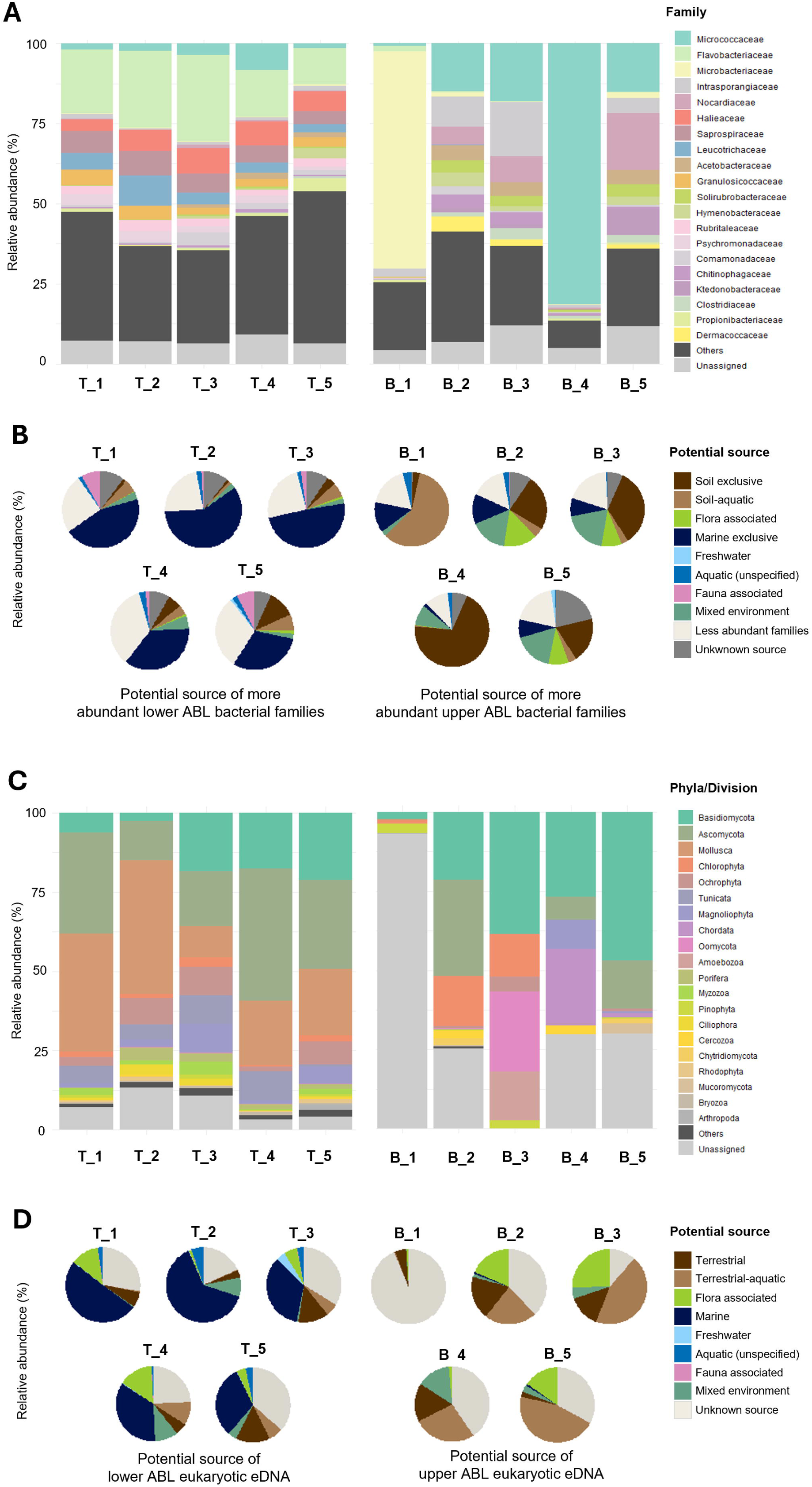
Microbial community composition in the lower and upper ABL layers. **A**. Relative abundance of the 20 most abundant airborne bacterial families across samples. *Others* includes less abundant families, while *Unassigned* includes uncultured families and taxa not assigned to any known family. **B**. Relative contribution of potential source ecosystems for the most abundant bacterial families. Families were considered abundant if they reached >1% relative abundance in at least one upper ABL sample (n = 32 families) or in at least one lower ABL sample (n = 35 families), evaluated independently for each height group (see supplementary Fig. S4 for the complete list of abundant families for each ABL layer). *Less abundant families* correspond to those with <1% relative abundance for which source information was not compiled from the literature. *Unknown source* represents families whose genera could not be assigned to any ecosystem or that include uncultured taxa lacking ecological context. See Supplementary Dataset S4 and S5 for bacterial genus-level literature-based source ecosystem assignments. **C**. Relative abundance of the 20 most abundant eukaryotic phyla/division across samples. *Other* includes less abundant phyla/division, while *Unassigned* includes uncultured phyla/division and taxa not assigned to any known group. **D**. Relative contribution of potential source ecosystems considering all detected eukaryotic taxa. *Unknown source* represents taxa which could not be assigned to any ecosystem or that include uncultured taxa lacking ecological context. See Supplementary Dataset S7 for eukaryotic genus-level literature-based source ecosystem assignments. **A-D.** Samples labelled T_1-T_5 correspond to the lower ABL samples, whereas B_1-B_5 correspond to the upper ABL samples.

Eukaryotic communities also displayed marked vertical stratification across the ABL (Fig. 2C). Four kingdoms were detected, with higher richness in the lower ABL, and all showed contrasting distributions between layers (Dataset S7). A substantial fraction of eukaryotic sequences could not be confidently assigned to any kingdom, particularly in the upper ABL (Dataset S7). Lower ABL communities were dominated by marine Mollusca, followed by Ascomycota and Basidiomycota, as well as Magnoliophyta and Ochrophyta, which exhibited relatively similar taxonomic profile across sampling events (Fig. 2C-D; Fig. S5; Dataset S7). In contrast, upper ABL communities were more variable but consistently dominated by Basidiomycota (Fig. 2C; Dataset S7) and enriched in soil-, flora- and freshwater-associated lineages (Fig. 2D; Fig. S5)

Building on these broad taxonomic patterns, a closer examination of the distribution of individual bacterial families revealed pronounced asymmetries in exclusivity across the ABL. Lower ABL communities contained many exclusive families (191 of 311), accounting for 15.1-24.4% of the community (Fig. S6A). Two exceeded 5% in at least one lower ABL sample (Fig. 3A): Streptosporangiales Incertae Sedis and Gottschalkiaceae, associated with Antarctic soil and fauna, respectively (Dataset S6). Only the latter was a significant indicator family of low altitude (Fig. 3A). The remaining exclusive families (189) contributed < 2% per sample, except Psycrhomonadaceae, represented by a psychrophilic marine genus (Dataset S6). Despite their low abundance, 37 of these 189 families were significant indicator families of low altitude, dominated by marine genera and representatives associated with Antarctic fauna (Dataset S6; Fig. S6A). In contrast, upper layer communities contained fewer exclusive families (14 of 134) contributing 0.6-8.4% (Fig. S6B), with Ktedonobacteraceae as the only significant indicator family of high altitude (Fig. 3A). This family was represented by one genera associated with pristine soils and cryospheric environments (Dataset S4).

**Fig. 3.**
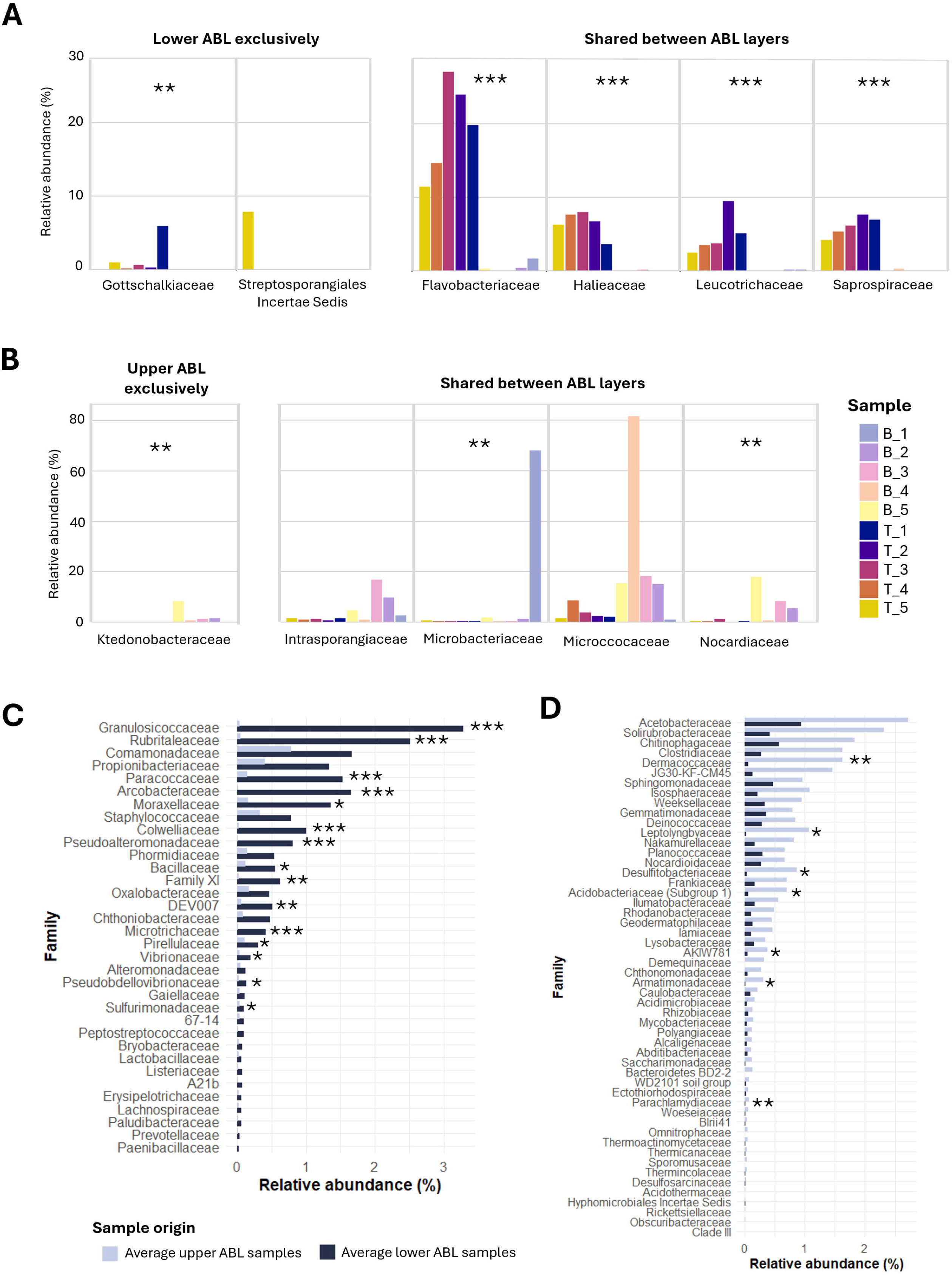
Representative bacterial families associated with the lower and upper ABL layers. **A.** Families representative of lower ABL layer exhibiting >5% difference in at least one sample compared with their corresponding upper counterpart. Both families exclusive to lower ABL layer and those shared between layers that meet this criterion are shown. **B.** Families representative of upper ABL layer exhibiting >5% difference in at least one sample compared with their corresponding lower counterpart. Both families exclusive to upper ABL layer and those shared between layers that meet this criterion are shown. **A-B**. Samples labelled T_1-T_5 correspond to the lower ABL samples, whereas B_1-B_5 correspond to the upper ABL samples. **C-D**. Families with <5% difference between layers but classified as lower-layer (C) and upper-layer (D) representative families based on average abundance: families are considered lower-layer representative taxa when their average abundance at low elevation is at least twice that at high elevation, and vice versa for high elevation. **A-D**. Significantly indicator families based on the IndVal test are marked as follows: *** (*p* = 0.001), ** (*p* < 0.01), * (*p* < 0.05).

At eukaryotic level, a similar but weaker pattern was observed, with a higher number of exclusive orders in the lower ABL (Fig. S7). Five orders exceeded 5% difference in at least one sample in the lower ABL, including three marine orders within Animalia (Myoida, Veneroida and Stolidobranchia) and two fungal flora-associated Lecanorales and Thelebolales orders (Dataset S7). All except Thelebolales were significant indicator orders of low altitude (Fig. S7A). In the upper ABL, only three exclusive orders exceeded a 5% difference, although none were significant indicator orders: the protist terrestrial-aquatic order Arcellinida and Peronosporales, and one fungal order Candelariales linked to flora-associated taxa not reported in Antarctic reference records (Fig. S7B; Dataset S7). Lower-layer exclusive orders below the 5% threshold contributed 8.7–22.68%, whereas upper-layer exclusives contributed only 0–1.42% (Fig. S7).

Beyond exclusivity, several bacterial families shared across ABL showed consistent height preferences. Eight families differed by more than 5% in relative abundance between layers in at least one sampling event. These lower-layer representative families were defined by Gram-negative families included the marine-associated Flavobacteriaceae, Halieaceae, Leucothrichaceae and Saprospiraceae, all significant indicator families of low altitude composition (Fig. 3A; Dataset S6). In contrast, Intrasporangiaceae, Microbacteriaceae, Micrococcaceae and Nocardiaceae, Gram-positive families were representative of the upper-layer and associated with cryospheric soils (Fig. 3B; Dataset S4), with Microbacteriaceae and Nocardiaceae identified as significant indicator families of high altitude (Fig. 3A). Bacterial families showing less than 5% differences also exhibited directional trends when averaged across events. Using a twofold difference in mean relative abundance between atmospheric layers as a threshold, 34 families showed preference for the lower ABL, 15 of them were significant indicator families (Fig. 3C). These lower-layer representative taxa are associated primarily with marine genera, although some were associated with soil and freshwater (Dataset S6). Conversely, 52 bacterial families showed high-altitude preference, including seven significant indicator families (Fig. 3D), and are associated with cryospheric soil genera (Dataset S4). Finally, 26 families showed no height preference, all of them with low abundance in both layers, and none were identified as significant indicator families (Fig. S8).

Eukaryotic orders also showed height-associated patterns, although in fewer taxa. Seven fungal orders frequently detected in Antarctica were identified as representative taxa in the upper ABL (Helotiales, Polyporales, Leucosporidiales, Sporidiobolales, Cystofilobasidiales, Filobasidiales and Tremellales), whereas Pleosporales (terrestrial-aquatic fungi) and Syndiales (marine protist) were representative taxa of the lower layer (Dataset S7). Indicator analysis detected no significant upper-layer indicator orders, while nine orders were significantly associated with the lower ABL (Table S1). Together, these results support a strong vertical stratification of microbial taxa within the ABL, generating distinct airborne assemblages at each height both prokaryotic and eukaryotic domains.

In terms of microbial abundance, the cell concentration average in the upper ABL was significantly higher than in the lower ABL (7.68 × 10^3^ vs. 5.41 × 10^2^ cells m^−3^), with sampling event 5 in the upper ABL reaching 3.54 × 10^4^ cells m^−3^ (Fig. 4A). Across both layers, the microbial community was dominated by particles smaller than 5 µm, accounting for over 90% of detected events in all samples (Fig. 4B, 5A). Morphological differences were observed between the lower and upper ABL. While coccoid forms were predominant at both heights, filamentous morphologies were more prevalent in the upper ABL than the lower ABL (Fig. 4B). Intermediate-sized particles (5–20 µm) were more frequently detected in the lower ABL than in the upper ABL, although not significantly (Fig. 4B). Large particles (>20 µm) were rare at both altitudes with no significant differences (Fig. 4B). These larger particles resembled filamentous cyanobacteria and eukaryotic algae (Fig. 5B-C), diatoms (Fig. 5D) and pollen (Fig. 5E). During sampling event 5, structures compatible with fungal forms were detected at upper ABL, a feature not observed in other events (Fig. 5F). Most <5 µm particles appeared as free cells or in pairs (Fig. 5A,G), or in small aggregates (Fig. 5G) rather than attached to non-biological larger particles (Fig. 5H). Samples from the lower ABL had more non-biological debris and fibres, adding background noise compared to the upper ABL. Colonies of bacilli and cocci were observed at both altitudes (Fig. 5I).

**Fig. 4.**
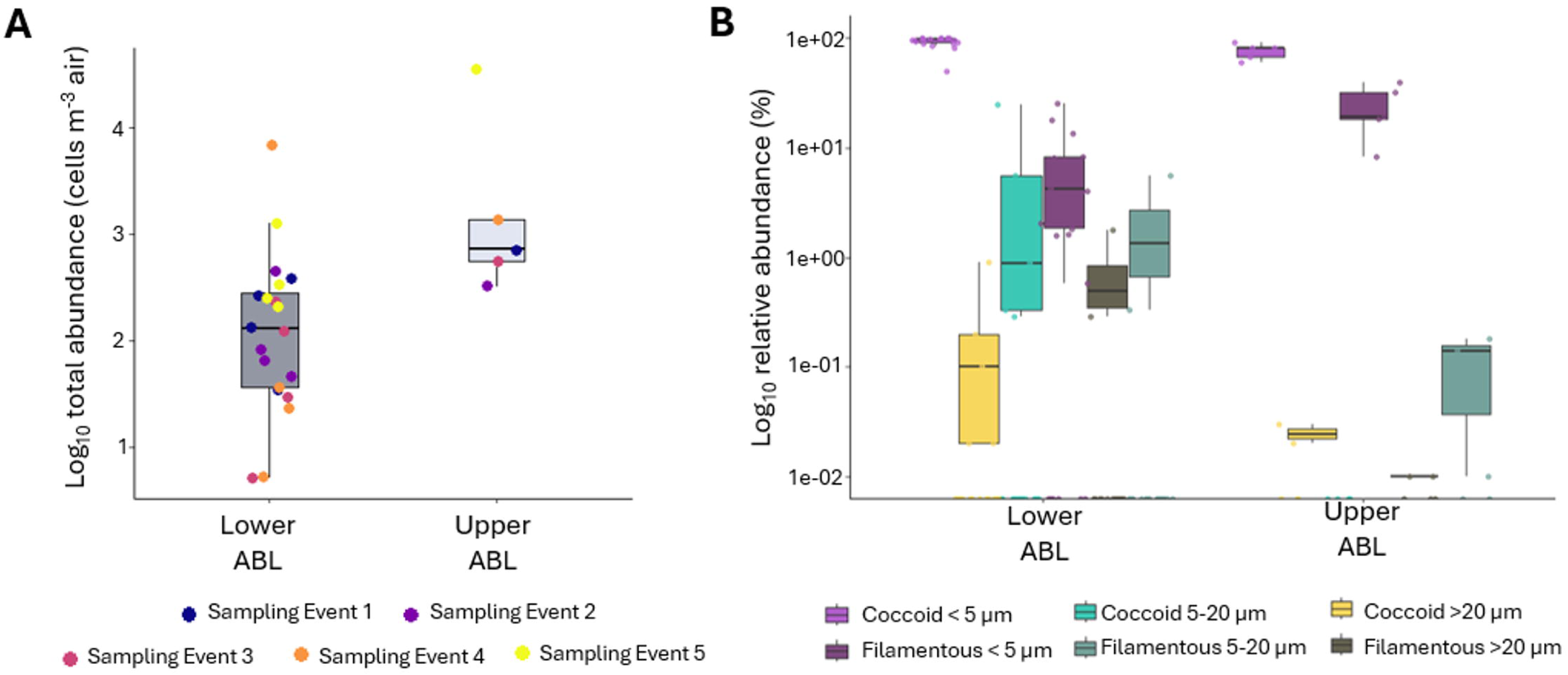
Airborne microbial abundance and morphological composition in the lower and upper ABL. **A.** Total airborne microbial abundance recorded for each sample in the lower and upper ABL. Average abundance differs significantly between layers (two-way ANOVA, *p* = 0.01), whereas the effect of sampling event was not significant (*p >* 0.05). **B.** Relative abundance of particle size classes and morphologies in the lower and upper ABL. The relative contribution of coccoid and filamentous particles < 5 µm differed significantly between layers (Wilcoxon signed-rank test, *p* < 0.05). No significant differences were observed for the remaining categories (Wilcoxon signed-rank test, *p* > 0.05).

**Fig. 5.**
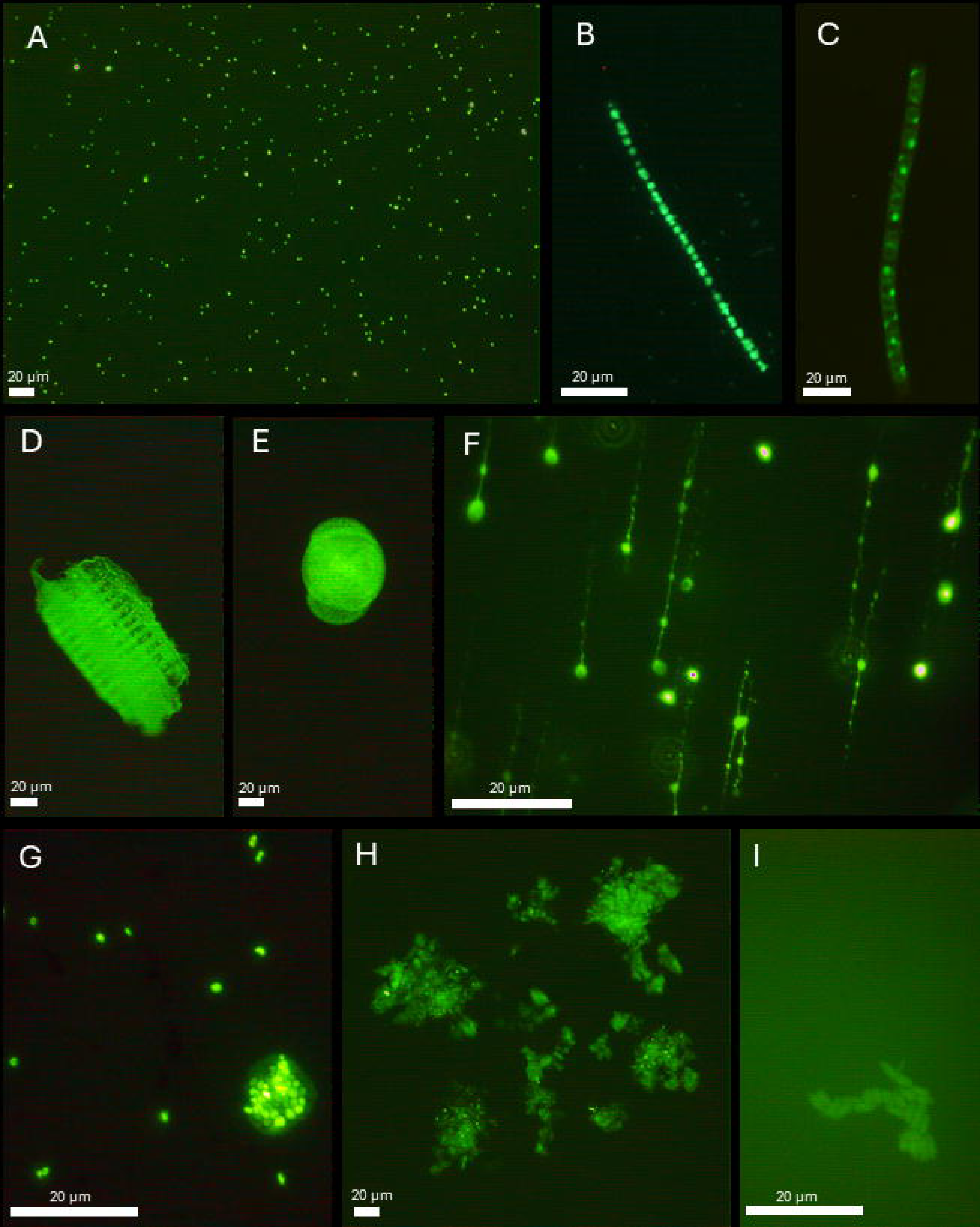
Representative airborne microbial particles detected in the lower and upper ABL. Airborne microorganisms were observed by epifluorescence microscopy at 1000× magnification using SYBR Green I as fluorochrome. **A.** Representative fields showing the dominance of particles < 5 µm in both ABL layers. **B.** Filamentous cyanobacteria. **C.** Filamentous eukaryotic algae. **D.** Structure compatible with a diatom. **E.** Structure compatible with a pollen. **F.** Structures compatible with fungal forms detected during sampling event 5 at both altitudes. **G.** Free individual < 5 µm cells, pairs and small aggregate. **H.** Non-biological particles to which microbial cells were occasionally attached. **I.** Colonies of bacilli observed at both the lower and upper ABL.

### Differences among microbial communities in the upper ABL

Microbial communities in the upper ABL exhibited pronounced variability among the samples, contrasting with the relative temporal homogeneity observed in the lower ABL. The strongest differences were observed during events 1 and 4, which shared relatively few bacterial ASVs and families with the remaining samples (although they represented only a minor fraction of the community) and displayed distinctive community structures (Fig. 6A, Dataset S3). Event 1 was characterized by a community largely associated with marine environments, whereas event 4 showed evidence of a stronger terrestrial influence. Notably, the eukaryotic community composition of the latest displayed the highest relative abundance of Animalia among all events, mainly represented by Aves- and Mammalia-associated sequences (Dataset S7). These biological differences were consistent with the atmospheric transport patterns inferred from back-trajectories and LCS analyses (Fig. 6B). Event 1 was associated with predominantly marine air masses and limited northward transport across the Antarctic Convergence. In contrast, air masses during event 4 crossed Livingston Island at low altitude (data not shown) and LCS analyses showed relatively isolated convergent flows primarily connected to the Antarctic Peninsula.

**Fig. 6.**
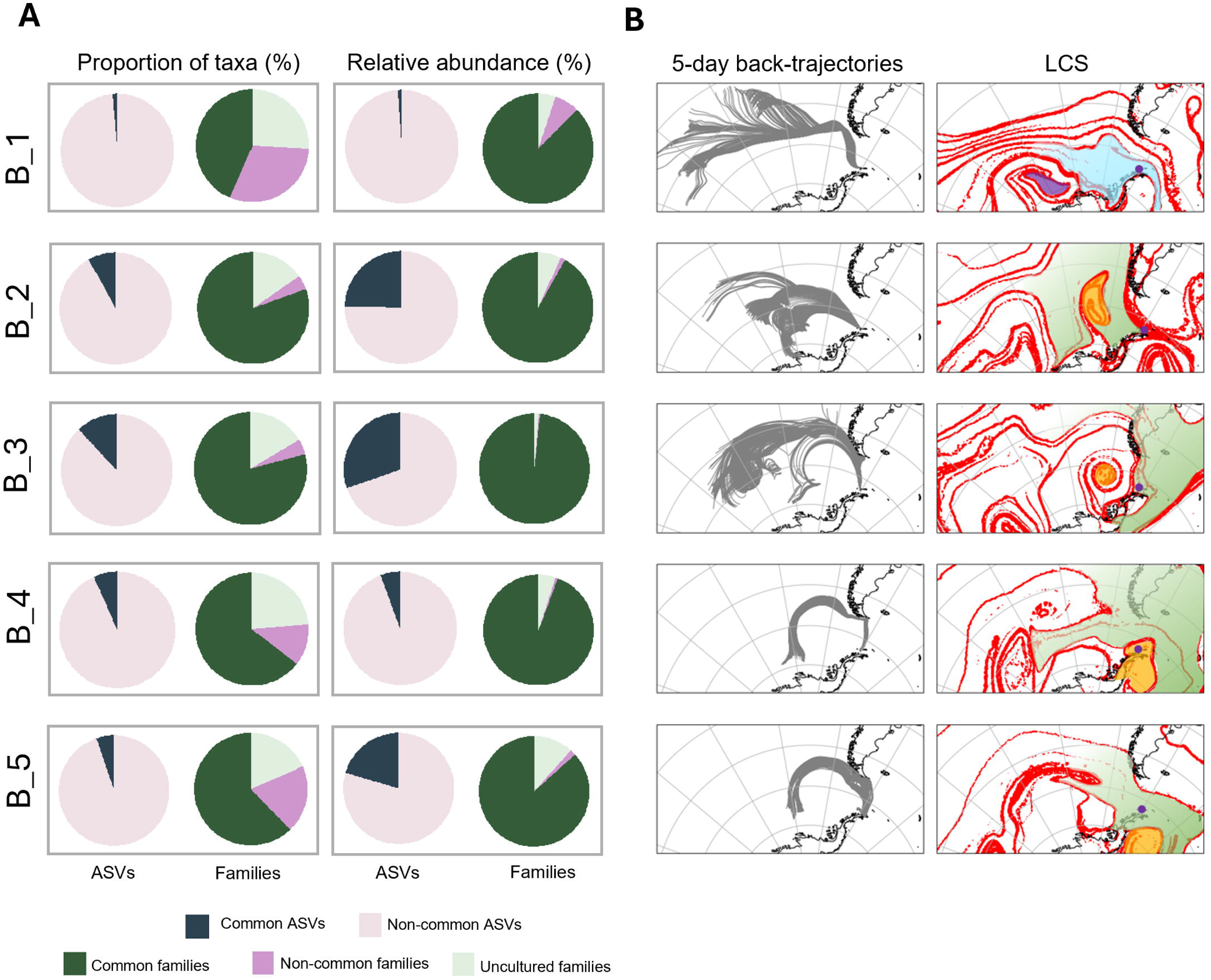
Similarities and differences among upper ABL bacterial communities and associated air mass trajectories. **A.** Proportions of shared and exclusive taxa at the ASV and family levels across upper ABL samples. Taxa were considered shared when detected in at least one additional upper-layer sample. The fraction of ASVs/families in each sample that were shared or exclusive, together with their relative contribution to community composition, is shown. **B.** Five-day back-trajectories of air masses associated with upper ABL samples and associated attracting Lagrangian coherent structures (LCS). Colors of main air masses are added to help tracking. Note that trajectories without dry deposition are shown as they are representative of those obtained when applying dry deposition for the most abundant morphotypes which represent the observed microbial community (see Fig. S9).

The remaining events showed broadly similar bacterial compositions and potential terrestrial source environments (Fig. 6A; Dataset S4). Although eukaryotic communities exhibited some variation in the relative abundance of major groups, with event 3 enriched in protist and event 5 characterized by a higher fungal contribution and lower representation of Plantae, their potential source environments remained broadly consistent across events (Fig. 1D, Dataset S7). For these events, trajectory analyses indicated either transport from southern South America or long-range marine transport, In agreement LCS patterns revealed a north-south atmospheric connection between South America and the sampling location (Fig. 6B). Together, these results suggest that temporal variability in the upper-ABL microbial communities was largely driven by changes in air-mass origin and transport history, which modulated the relative contribution of marine and terrestrial sources.

## DISCUSSION

Our study demonstrates a clear vertical stratification of airborne microbial communities across the Southern Ocean ABL. The results revealed two sublayers with contrasting microbial assemblages and dynamics in both bacterial and eukaryotic communities across taxonomic resolutions. This vertical organization could reflect limited vertical mixing within the regional MABL, which is frequently decoupled into a surface layer and an overlying mixed layer [21,41]. Under these conditions, particles emitted near the surface tend to remain confined to lower atmospheric levels [42], whereas episodic disturbances such as cyclonic systems can enhance upward transport into higher layers [21,41]. In contrast, Kobayashi (2022, 2026) [14,15] reported relatively similar bacterial community composition between near the ground and upper air layers in coastal East Antarctica, which may reflect the stronger influence of katabatic winds and coastal hydraulic jumps that promote vertical mixing within the boundary layer in that region.

In the lower ABL, microbial communities were similar along the sampling period, dominated by taxa commonly reported in Antarctic Peninsula near-surface studies, both bacteria [43–45] and eukaryote [46–48]. These communities closely resembled nearby environmental matrices, a pattern interpreted as indicating a predominant local influence [45,49]. The diverse landscape of Byers Peninsula, encompassing marine and terrestrial environments, as well as animal colonies [22], likely acts as a mosaic of microbial sources. Although marine aerosolization is considered to contribute less microbial biomass to the atmosphere than terrestrial sources [50], the strong marine signal observed in our results reflects the influence of the Southern Ocean, potentially amplified by the region’s limited terrestrial area. Bacteria associated with animal microbiomes supports inputs from abundant seabird and pinniped colonies in this region [45,51], while eukaryotic communities include taxa linked to characteristic vegetation, particularly mosses and lichens widespread across the peninsula [52]. The limited variability in community composition among samples reflects efficient turbulent mixing of microorganisms from multiple local sources within this surface layer [21,41] under comparable meteorological conditions (Dataset S2) [49, 53]. Together, these findings suggest the lower ABL is a well-mixed microbial reservoir reflecting surrounding ecosystems.

In contrast, microbial communities at the upper ABL-free troposphere interface showed higher variability among samples, which is consistent with the influence of air mass history and long-range atmospheric transport [3,5]. At these heights, microbial assemblages integrate inputs from regional and continental sources, according to air mass back-trajectories [44,54,55], while their dispersal may be further constrained by Lagrangian transport barriers [56,57]. This is illustrated by the shifts in community composition associated with changes in the horizontally advected air masses (Fig. 6B). The similarities between samples 3 and 5, despite the transient change observed in sample 4 during the passage of a different air mass, suggests that microbial communities are relatively homogeneous within a given air mass and that atmospheric transport barriers can also act as effective barriers to microbial communities at high altitude.

Despite clear compositional differences between atmospheric sublayers, our results indicate vertical connectivity within the air column. Although only a small fraction of ASVs was shared, most bacterial families and eukaryotic orders detected aloft were also present in the lower layer, whereas many lower-ABL taxa were absent at higher altitudes. While the observed ASV overlap could reflect bidirectional exchange between atmospheric layers (i.e., upwelling and downwelling), the higher-level taxonomic asymmetry is consistent with a selective upward transport [58], whereby only a subset of surface-derived microorganisms is transferred to upper atmospheric levels within the stratified Southern Ocean MABL [41,42]. A fraction of this filtering appears to depend on both potential source and particle characteristics, with a shift from marine-associated taxa in the lower ABL to soil- and flora-associated lineages aloft, accompanied by a higher abundance of larger and aggregated particles in the lower ABL. Marine-associated microorganisms, potentially transported via sea-spray droplets, and aggregates exhibit larger effective size and density, reducing vertical transport efficiency, whereas drier and discrete cells may be more aerodynamically stable [32,58]. Similar patterns have been reported for non-biological aerosols in the Southern Ocean MABL, where sea-spray droplets dominate lower ABL and drier sea-salt particles prevail aloft [59]. Consistently, the <5Oµm fraction contained a higher proportion of linear forms in the upper layer, a morphology associated with enhanced transport potential [32]. Such constraints may disproportionately affect larger eukaryotic cells and spores relative to bacterial cells, potentially explaining the greater variability of upper-layer eukaryotic assemblages in upper-ABL events despite having similar air-mass trajectories and bacterial communities. This eukaryotic pattern may be further reinforced by inversion layers that restrict exchange near the ABL–free troposphere transition.

In addition to physical transport constraints, microorganisms reaching higher atmospheric layers are exposed to intense UV radiation, desiccation, oxidative stress and large temperature fluctuations, all of which can contribute to the degradation of cellular structures and associated genetic material [1,3,58]. Accordingly, altitude may act as a filtering process that reduces the diversity of detectable genetic material, resulting in upper-ABL communities that represent a subset of those observed in the lower ABL [58]. Whether these upper-ABL taxa are active or viable is not known and their presence may arise from differential preservation of biological particles and DNA during atmospheric transport. Consistent with this interpretation, several taxa enriched in the upper layer possess traits that may enhance the preservation of cellular integrity under atmospheric stressors. These included Gram-positive lineages which are more resilient to desiccation and UV-induced DNA damage [60]. In contrast, Gram-negative bacteria, were more abundant in the lower layer. They may interact electrostatically with larger particulate matter due to their surface properties, potentially limiting their vertical transport [3,50]. Upper-ABL indicator bacterial taxa included psychrophilic and psychrotolerant genera (Dataset S4) within Microbacteriaceae (*Cryobacterium* and *Parafrigobacterium*), Micrococcaceae (*Specibacter*), Chitinophagaceae (four psychrophilic genera) and Desulfitobacteriaceae (*Desulfosporosinus*). Additional enriched lineages, including Nocardiaceae (*Nocardia* and *Rhodococcus*) and Intrasporangiaceae (*Knoellia*), possess traits associated with resistance to desiccation, UV exposure, and DNA damage [61–63], while pigmented members of Micrococcaceae and Deinococcaceae may be less susceptible to photodamage [1,58]. Likewise, globally distributed airborne genera such as *Sphingomonas*, *Hymenobacter*, and *Pseudomonas* were also abundant in the upper layer [64]. These bacterial taxa possess traits favouring persistence in atmospheric environments and therefore contributing to the detectability of biological particles and their genetic material under prolonged atmospheric exposure [1]. Beyond bacteria, the atmosphere also transports a wide range of eukaryotic propagules, including fungal and plant spores, pollen grains, and small seeds, with protective structures adapted for wind dispersal and preservation of cellular and genetic integrity during transport [65]. However, detected eukaryotic DNA may also derive from fungal hyphae, plant tissues, or other fragmented biological material [66], as suggested by microscopic observations in this study. Several fungal indicators of the upper layer belonged to groups associated with cold environments (Dataset S7) and photoprotective traits [67], including *Rhodotorula*, *Cystofilobasidium*, *Mrakia*, Bulleribasidiaceae, and *Filobasidium*. Similarly, taxa such as the lichen-associated *Chloroidium* [66,68] and snow alga *Chloromonas* [69] are characteristic of environments exposed to intense radiation, desiccation, and low temperatures. Together, these observations support the idea that altitude acts as a filtering environment in which only a subset of the biological material aerosolized near the surface remains detectable at higher altitudes. The enrichment of particular bacterial and fungal lineages in the upper ABL is therefore consistent with differential cellular preservation and associated genetic material under harsh tropospheric conditions, contributing to the lower alpha diversity observed in the higher layer.

Microbial concentrations in the upper ABL were higher than in lower ABL, suggesting that this signal may progressively enrich for biological particles, in contrast to lower-ABL communities which are continuously reshaped by deposition and resuspension processes. Environmental conditions in the Southern Ocean MABL may further enhance this persistence, as multilayer mixed-phase cloud systems frequently occur at low altitudes ∼500 m during summer [21,41]. These clouds can act as transient “oases” for those potentially active microorganisms in an otherwise dry, nutrient-poor atmosphere, providing liquid water and dissolved organic nutrients while partially shielding microorganisms from UV radiation [3,70]. Conversely, bioaerosols can influence cloud microphysics. Microorganisms and pollen act as efficient ice-nucleating particles (INPs), initiating ice formation at warmer subzero temperatures than most inorganic particles [3,66]. Several bacterial and eukaryotic genera detected in our upper ABL samples have previously been associated with ice-nucleation activity [66]. Enrichment of plant-associated taxa has also been reported in cloud and hailstone microbiomes [71], and these taxa were the dominant component of our upper atmospheric samples. During sampling at 1,000 m (event 5), the balloon ascended above a cloud layer (Video S1-2), and ice fragments were observed on the tether line (Fig. S10). Filamentous structures resembling fungal hyphae (Fig. 5F), consistent with *Mortierella* [72], were detected, together with *Leucosporidium*, highly abundant in this event and known for ice-nucleation activity [73,74]. This event also showed the highest microbial abundance. Although anecdotal, these observations are consistent with clouds acting as transient reservoirs of airborne microorganisms with potential microphysical interactions.

Microorganisms remaining in an accumulation mode in the upper ABL may contribute to long-range atmospheric dispersal, as reflected by upper-layer-exclusive taxa such as Ktedonobacteraceae. This family has been reported in nutrient-poor environments, including deserts and cold regions [75], and all ASVs show >98% sequence identity with sequences from South America and New Zealand (Dataset S8), supporting their dispersal potential. This pattern is consistent with west-to-east atmospheric circulation over Antarctica and with the idea that particles reaching the free troposphere, where airflow is faster and more uniform, can be rapidly transported over large distances [60]. Their dominance at high altitude, especially during sampling at 1,000 m (free troposphere), suggests prolonged atmospheric residence with potential contribution to colonization of nutrient-poor early-stage soils in distant regions should them remain viable [76]. However, long-range atmospheric dispersal is not restricted to high altitudes. Although near-surface airborne microbial communities often reflect local emission sources, some sampling events showed greater similarity in ASV composition between lower and upper layers when air masses at low ABL shared similar back-trajectories with upper ABL (Supplementary Results; Fig. S9, S11), supporting the history of air-mass movements can influence microbial composition at different altitudes.

Overall, these results provide a representative overview of the vertical distribution of airborne microorganisms and propagules in coastal polar environments with low topographic complexity and strong marine influence, typical of the Southern Ocean during the austral summer. Our data support a conceptual framework in which microbial community structure within the ABL is organized into two partially decoupled sublayers. The lower surface layer is dominated by turbulent mixing and strong local inputs, resulting in relatively homogeneous microbial communities, although contributions from long-range transport are also detectable. In contrast, the upper layer is shaped by long-range transport and environmental filtering, favouring the presence of microorganisms capable of withstanding atmospheric stress and longer residence times. This vertical heterogeneity is consistent across taxonomic scales: ASV-level differences reflecting short-term processes such as vertical mixing and horizontal transport, whereas family-level patterns indicate broader ecological affiliations and selective pressures influencing persistence across atmospheric heights over longer timescales. These findings highlight the ABL as a structured environment rather than a passive transport medium, where physical dynamics and biological characteristics jointly determine microbial retention and dispersal.

Ice-free coastal areas of the South Shetland Islands, characterized by low altitude and simple topography, provide an ideal natural laboratory for investigating long-range atmospheric transport. Their geographic position as one of the first Antarctic regions reached by air masses carrying terrestrial particles from South America [60, 77] facilitates the study of long-range dispersal while minimizing the influence of surrounding terrestrial sources. Moreover, the Antarctic Peninsula coastal zone is especially sensitive to environmental variability, as its strong maritime exposure and reduced glacial buffering make it among the first Antarctic regions to respond to changing conditions [78, 79]. This combination of climatic sensitivity and connectivity makes these systems particularly relevant for understanding how shifts in atmospheric dynamics may alter microbial dispersal pathways and exchange. However, this framework does not apply uniformly across Antarctic coastal systems. Regions with complex topography, such as the McMurdo Dry Valleys, exhibit terrain-driven atmospheric dynamics that may disrupt ABL homogeneity and alter vertical exchange through slope flows and localized circulation patterns [20,80], as well as katabatic winds [81]. In addition, diurnal surface heating of exposed soils can further modulate ABL structure [82]. In contrast, marine-influenced systems such as the South Shetland Islands show more stable ABL conditions due to oceanic thermal inertia. Finally, the necessarily fair-weather conditions needed during samplings to allow the use of the tethered ballon limit our assessment of the influence of episodic events such as precipitation or strong winds. Future research should integrate longer-term observations, atmospheric transport modelling, and functional characterization of airborne microorganisms to better resolve microbial responses across atmospheric regimes and improve understanding of how atmospheric processes shape microbial dispersal, ecosystem connectivity, and biogeographic patterns in polar regions.

## ACKNOWLEDGEMENTS

The authors are grateful to Cristina Casero, Andrés Alastuey, Luis Piñuel, José Antonio López Orozco, Paula Martínez, Pilar González-García and Sara Granados for their valuable recommendations and assistance in the preparation of sampling materials.. We also thank the staff at SEGAINVEX-UAM, Calypso Instruments and Smarthaps Company, especially Nuria Diaz, Juan Ramón Marijuán, and Francisco Navarrete. The authors are grateful to the Unidad de Tecnología Marina (UTM-CSIC), the crews of the BIO Hespérides (Spanish Navy), and AEMET-Antarctica group for logistical and meteorological support during Antarctic campaigns. Special thanks are extended to Benito Elvira and Hilo Moreno. The authors acknowledge the computational resources, technical expertise, and assistance provided by the Centro de Computación Científica at the Universidad Autónoma de Madrid (CCC-UAM), as well as the NOAA Air Resources Laboratory (ARL) for providing the HYSPLIT transport and dispersion model.

## AUTHOR CONTRIBUTIONS

**S.G.** carried out conceptualization, methodology, investigation, data curation, formal analysis, visualization, writing the original draft, and writing review and editing. **W.Y.K.** performed methodology and writing review and editing. **P.S.** carried out methodology and writing review and editing. **T.P.** performed methodology, formal analysis, software, visualization, and writing review and editing. **J.A.H.** carried out methodology and writing review and editing. **M.B.** carried out conceptualization, methodology and writing review and editing. **J.M.** carried out methodology and writing review and editing. **K.A.** carried out methodology. **S.G-H.** performed conceptualization, methodology, investigation, formal analysis, software, visualization, writing the original draft and writing review and editing. **A.J.** was responsible for conceptualization, methodology, investigation, formal analysis, software, funding acquisition, project administration, resources, supervision and writing review and editing. **A.Q**. was responsible for conceptualization, methodology, investigation, funding acquisition, project administration, resources, supervision and writing review and editing.

## DATA AVAILABILITY

The raw DNA of bacteria and eukaryotic sequences generated in this study have been deposited in the NCBI Sequence Read Archives under BioProject accession number PRJNA1484524 and PRJNA1484830, respectively. All data used in this study, including ASV abundance tables, associated sequences, taxonomic assignments, and physical datasets, are available at Zenodo (https://doi.org/10.5281/zenodo.21217775). The physical datasets include balloon micrometeorological measurements and flight height calculated at 1-min intervals for each sampling event, the original back-trajectories and the corresponding back-trajectories with dry deposition applied, as well as the LCS datasets for each sampling event. Five-day back trajectories were computed using HYSPLIT. All sampling coordinates and times are provided in Dataset S1 to ensure full reproducibility.

## STUDY FUNDING

This work is part of the MICROAIRPOLAR-2 project (PID2020-116520RB-100), funded by MCIN/AEI/10.13039/501100011033. Sofía Galbán was supported by a PIPF-contract fellowship (PIPF-2022/ECO-25833) from Comunidad Autónoma de Madrid government’s (Spain). This work was supported by PEJ-contract fellowship (PEJ-2024-AI/COM-32565) financed by Comunidad Autónoma de Madrid government’s (Spain) and co-financed by European Union. Tamara Pletzer was supported by the New Zealand Ministry of Business, Innovation and Employment through the Antarctic Science Platform (ANTARCTICANZ2504).

## Supplementary Methods

### Sampling

Airborne microbial communities were captured using a custom-designed collector system optimized for polar conditions and minimal human intervention. For full design specifications, see Parro et al. (2025) [23]. Each collector housed a removable sterile support coated with Vaseline® over a 70.68 cm² surface. The collectors were modified from that of Parro et al. (2025) [23] as follows: shortened to reduce weight in the balloon and a tail fin was added to ensure passive alignment of the inlet with the wind direction (Fig. S1). Before each sampling, collectors were run for 5 min without the Vaseline-coated support at maximum speed, with the operator positioned downwind. Sterile supports were then inserted under aseptic field conditions (gloves and face mask), and collectors were raised to target height and remotely activated. Sampling was performed simultaneously at both platforms (tower and balloon) for 6 h, passing 275.4 m^3^ of air. After sampling, the collectors were remotely stopped and, in the case of balloon recovered at constant speed with a homemade electronically controlled winch. The supports were retrieved into sterile Whirl-Pak® bags, using gloves and face masks, and stored with negative controls at ≤ -20°C until processing in UAM laboratories. Negative controls included: (i) an unopened Vaseline-coated support transported to the field to verify sterility during preparation and transport (CNN), and (ii) a Vaseline-coated support exposed but non-active collectors at each height after decontamination (CNS) to verify the effectiveness of the collector decontamination procedure.

For upper ABL sampling, the collector was installed in an instrumented sampling gondola suspended 6 m below a spherical tethered balloon (Fig. S1). The tethered balloon system was specifically designed for high-altitude profiling, capable of carrying instruments up to 1200 m above ground level [83]. Besides the biological collector, the gondola equipped with a commercial ultrasonic anemometer (Calypso ®) and temperature, humidity and pressure sensor (Disc Maxi 4-in1, BlueMaestro ®) that recorded data at 1 min intervals, and a GoPro camera. Sampling was conducted only under wind speeds < 25 km/h for safety and operational stability of the balloon. Balloon height (Fig. S3) was calculated with the formula h=8.5 x (P_surface_ - P_balloon_), where *P_surface_* and *P_balloon_* are the surface and onboard barometric pressures, respectively. For the first flight, due to a sensor failure, balloon height was reconstructed using a quadratic model relating height with wind speed (Fig. S2), assuming that wind progressively displaces the tethered balloon from its maximum cable height.

To characterize the microbial communities at the lower ABL, a sampling tower was installed next to the balloon anchoring point. The tower was equipped with eight collectors, installed in pairs at four heights (7.5, 2.6, 0.6 and 0.3 m; Fig. S1). Surface pressure data was also recorded simultaneously with a temperature, humidity and pressure sensor (Disc Maxi 4-in-1, Blue Maestro®) placed at ground level.

### DNA extraction, amplification and sequence processing

Vaseline was recovered from the sample holders with a sterile blade, and the microbial cells were extracted in phosphate-buffered saline (PBS) with 0.005% Triton X-100, sterile-filtered (0.2 μm) and autoclaved at 110°C for 15 min. Of the 4 mL suspension obtained per sample, 200 μL were reserved for microscopy and the remaining volume used for genomic DNA extraction with the DNeasy PowerSoil Kit (QIAGEN)® following Parro et al. (2025) [23]. To ensure sufficient biomass for downstream analyses, samples from paired collectors at each tower height were pooled during DNA extraction (spin-column loading step) and during the filtration step for microscopy preparation. All procedures were performed under sterile conditions following best practises for low-biomass samples [31], including the use of personal protective equipment, laminar-flow hoods, surface decontamination with 4% sodium hypochlorite, and UV-C irradiation. Extracted DNA was stored at -20°C until amplification at Microomics Systems S.L. (Spain).

Bacterial community composition was characterized using primers 341F (5′-CCTACGGGNGGCWGCAG-3′) and 805R (5′-GACTACHVGGGTATCTAATCC-3′), amplifying V3-V4 hypervariable region of the 16S rRNA gene (∼464 bp) [24]. Eukaryotic communities were assessed using Earth Microbiome Project (EMP) primers 1391F (5′-TCGTCGGCAGCGTCAGATG TGTATAAGAGACAG GTACACACCGCCCGTC-3′) and EukBr (5′-GTCTCGTGGGCTCGGAGATGTGTATAAGAGACAGTGATC CTTCTGCAGGTTCACCTAC-3′) targeting the V9 hypervariable region of the 18S rRNA gene (120–260 bp) [25,26]. Libraries were sequenced on the Illumina MiSeq platform using paired-end sequencing of 300 cycles for bacteria and 150 cycles for eukaryotes. Field negative controls were processed alongside samples throughout extraction, amplification, and sequencing, together with positive and negative PCR controls. Second-PCR negative controls did not yield sufficient high-quality reads and were excluded from downstream analyses.

Demultiplexed pair end-reads, including controls, were processed with DADA2 [27]. Reads were first quality-filtered and trimmed to remove bases with a Phred score below 25. Amplicon sequence variants (ASVs) were inferred using error models for forward and reverse reads and later merged using default settings (≥12 nt overlap, 0 mismatches). Chimeric sequences and singleton were removed, as well as PhiX-derived reads, which represent a common Illumina sequencing artifact. Taxonomic assignment was performed against the SILVA v138.2 [28] using the ribosomal database project (RDP) naïve Bayesian classifier [29] implemented in DADA2 for bacteria, and the QIIME2 naïve Bayesian classifier [30] for eukaryotes. For eukaryotes, the classifier was previously trained on the target region using the corresponding primers. Chloroplast, mitochondrial, archaeal, and eukaryotic sequences were excluded from bacterial datasets, and chloroplast, mitochondrial, archaeal, bacterial, and kingdom-level unclassified sequences from eukaryotic datasets. To minimize contamination, all ASVs detected in field or PCR negative controls were conservatively excluded [31]. In the final curated dataset, 96.4 % of the total bacterial ASVs (96.9% reads) and 61.74% of the total eukaryotic ASVs (63.39% reads) were retained.

### Epifluorescence microscopy analysis of airborne microbial communities

For microbial abundance and morphometric analyses, 200 μL of each sampling cell suspension was stained with SYBR Green I (Thermo Fisher Scientific, Walthman, USA) to a final concentration 1x for 1h at room temperature in dark. Stained samples were then filtered onto black polycarbonate filters 0.2 μm pore size (Millipore) and mounted on microscope slides using EMOIL-F30CC (Olympus) as mounting medium.

Samples were examined using a Nikon Eclipse 80i epifluorescence microscope equipped with a mercury lamp and a long-pass FITC-LP01 filter cube (Semrock / IDEX-HS, part no. FL-007188) with a dichroic edge at 515Onm, transmitting 519–700Onm and efficiently blocking shorter wavelengths. This setup is optimized for SYBR Green I, which binds specifically to DNA and emits bright green fluorescence at ∼520 nn when excited at ∼497 nm, while remaining non-fluorescent when bound to other materials. Microbial abundance was estimated as the average count across 100 randomly selected fields of view at 1000x magnification. Cell counts accounted for Poisson-distributed variability to ensure overall precision (relative standard deviation <10%), even in low-biomass samples. Counted microorganisms were classified into three size categories (<5 μm, 5–20 μm, >20 μm) and two morphotypes within each category (coccoid and filamentous), following Galbán et al. (2021) [32]. Microbial lengths were measured using a Leica DFC300 FX camera and Leica Application Suite v.3.7.0. Final concentrations of airborne microbes were calculated as cells per cubic meter of air and the relative proportions of size and morphotype categories were expressed as percentages of the total counted cells. Differences in microbial abundance between lower and upper layers were tested using two-way ANOVA, whereas the Wilcoxon signed-rank test was used to compare relative contributions of size and morphotype categories between layers. Statistical significance was set at *p* < 0.05. Negative controls (CNN and CNS) consistently showed no detectable fluorescence, confirming the absence of contamination.

### Microbial community analyses

For all taxonomic composition analyses, samples from all tower heights were pooled by sampling event to obtain an integrated representation of near-surface airborne microorganisms within the first 10 m a.g.l. ASVs present in at least 60% of samples (Core60) were used to assess community consistency across sampling events within each layer for both bacterial and eukaryotic datasets. Exclusive and shared taxa were then identified for each ABL layer. Bacterial families and eukaryotic orders occurring only in one layer were considered exclusive, whereas those detected in at least one sample from both layers, regardless of sampling event, were classified as common to both layers and further examined to identify potential layer indicators. Each layer composition was also compared using the *layer-representative taxa*, defined using two complementary criteria. First, taxa showing a maximum difference in relative abundance greater than 5% between layers in at least one sampling event were included. Second, taxa not selected by the previous criterion were further evaluated only if they reached >1% relative abundance in at least one sample. These taxa were considered *layer-representative taxa* when their mean relative abundance in one layer was at least twofold higher than in the other layer. The layer-specificity of bacterial families and eukaryotic orders was further confirmed using the Indicator Species Analysis test (IndVal) with 999 permutations [37].

Differences in community composition among upper-layer sampling events were further explored using shared and exclusive ASVs, as well as bacterial families and eukaryotic orders. Taxa detected in more than one sampling event were considered shared, whereas those found in only one event were considered exclusive. For each sampling event, the proportion of shared and exclusive taxa relative to the total detected taxa for each sampling event community was calculated, together with their cumulative relative abundance. In addition, overlap between upper- and lower-layer communities within each sampling event was assessed by classifying bacterial ASVs and families as shared when detected in both layers and exclusive when restricted to a single layer.

### ABL height estimation

The ABL varies on timescales of hours to days. Its height can be estimated from the meteorological vertical profiles obtained during balloon ascent and descent (Fig. S3) using the Richardson number (*Ri*) in Eq. 1, assuming that each profile is measured quickly enough to avoid substantial changes in the ABL during the sampling period [38]. However, our measurements have some limitations, usual in all the profiles. When the denominator in Eq. 1 approaches zero (ΔU^2^ ≈ 0), *Ri* becomes artificially large and we therefore discarded *Ri* values computed where the wind-speed difference from the reference level was below 1.5 km h^−1^, the minimum difference we considered resolvable from noise. In addition, noisy data may arise from balloon movement, and to avoid this issue we calculate in practice the ABL height at the lowest altitude at which three consecutive data points exceed the *Ri* threshold. Because *Ri* is referenced to a fixed near-surface level, the height difference term in Eq. 1 grows continuously with altitude, so even weak background stratification can eventually push *Ri* above 0.25 well before the real capping inversion is reached. We therefore accepted a *Ri*-based crossing only when it coincided with enhanced static stability, requiring at least one of the three points to fall within the upper third of the Brunt-Väisälä frequency squared (N^2^) values of that profile. We considered the ABL height as the altitude of maximum N^2^ within that window. The sampling height ranges are shown in Table S2. In all cases except B_5, the ABL height was within the sampled height range, whereas B_5 was sampled exclusively within the free atmosphere.

### Atmospheric transport analysis

Five-day air mass back-trajectories were computed using the Hybrid Single-Particle Lagrangian Integrated Trajectory (HYSPLIT) model [39], a standard tool for reconstruction air-parcel pathways and transport patterns. Simulations were based on meteorological data from the global ERA5 system, with a 0.25 degrees resolution. Back-trajectories were initiated in the tower location and launched at one-minute intervals throughout the full sampling period. While the balloon trajectories were initialized at the estimated height each minute, the trajectories from the tower were initiated from an altitude corresponding to 0.1 of the ABL depth to capture representative conditions of the lower sublayer.

To better constrain the atmospheric transport potential of airborne microorganisms and account for the effects of particle size and morphology on air residence time, a dry deposition scheme was implemented following the methodology described in Galbán et al. (2021) [32] for both sampled upper- and lower-layer air masses. Particular attention was given to particles <5 µm, which are more representative of our samples as observed by microscopy. Spherical particles were defined based on measured diameters <5 µm across three observed size ranges (0.5, 2.5 and 4.5 µm). Linear particles were characterized using three observed equivalent spherical diameter (ESD) ranges (0.5, 1.5 and 2.45 µm). Simulations were conducted assuming 100% relative humidity and a particle density of 1 g cm^−3^. Back-trajectories calculated without dry deposition were found to be broadly representative of those including dry deposition (Fig. S9), indicating that the general back-trajectories analyses capture the potential maximum dispersion of the airborne microbial community dominated by particles <5 µm.

In addition, we computed the finite-time Lyapunov exponent (FTLE) to obtain the large-scale Lagrangian Coherent Structures (LCS) during the microbiological sampling period. FTLE analysis using backward-time integration provides a quantitative frame-invariant measure of the dominant attracting structures in the wind field [40]. These structures identify regions of air parcel converge that may influence airborne particle accumulation and transport. FTLE has been used to study atmospheric transport processes, including Saharan dust dispersion [84] and plant pathogen transport [57], as well as oceanic processes such as phytoplankton [85] and marine larvae aggregation [86]. For each sampling date, we performed 12-hour backward integrations from the end time of the sampling. This integration time was selected as a compromise between spatial resolution and noise reduction. Sensitivity analysis showed that longer integration periods blurred transport structures. We utilized ERA5 reanalysis data at the 925 hPa pressure level with 0.25° spatial resolution and NumbaCS Python package [87] for FTLE computation in atmospheric flows.

## Supplementary Results

### Similarities between lower and upper ABL layer bacterial communities across events

Despite the consistent differences observed between lower and upper ABL communities across sampling events, some events displayed partial similarities in bacterial composition between heights. In particular, events 2 and 3 showed a higher proportion of shared bacterial ASVs between layers compared to the remaining events (Fig. S11A-B). These shared ASVs were mainly assigned to bacterial families which were associated with upper ABL communities. Air mass back-trajectory analyses revealed similar transport pathways at both layers during these events (Fig. S9).

The strongest divergence between layers was observed during event 5, where the relative abundance of shared bacterial families between lower and upper-layer sample was lower than in all other events (Fig. S11B). This pattern coincided with contrasting air-mass back trajectories, with upper ABL samples linked to western Antarctic Peninsula and Southern Ocean origins, whereas lower ABL samples were associated with eastern pathways from the Weddell Sea (Fig. S9).

Overall, similarity in the bacterial communities at ASV level between two layers was associated with comparable air-mass transport histories, whereas divergent air-mass origins were reflected in differences in community composition at the family level. These patterns were particularly pronounced in communities from the upper ABL.

## Supplementary Figures

**Fig. S1.**
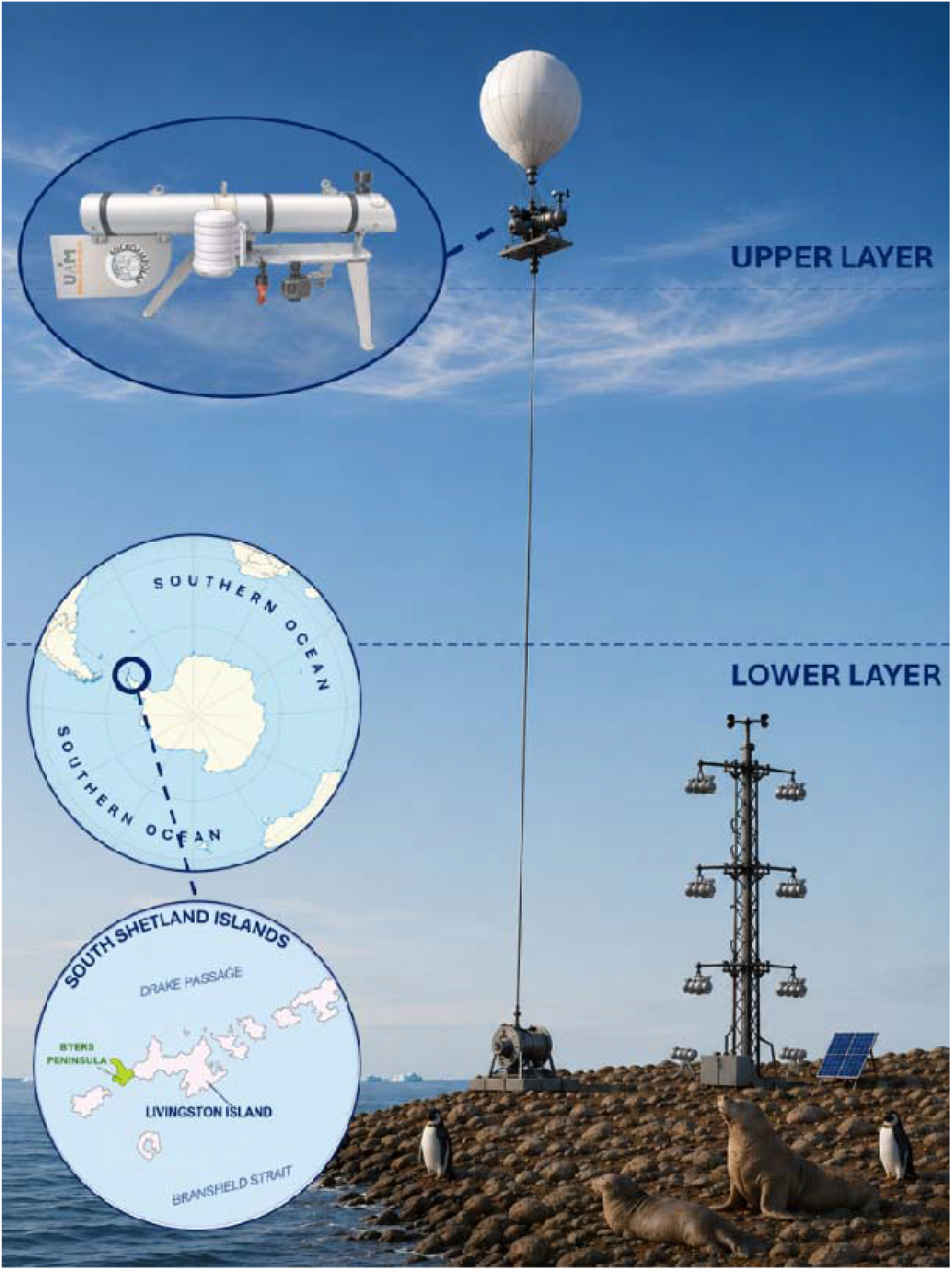
Overview of the vertical profile airborne microbial sampling design in the ABL. Airborne microbial communities were collected simultaneously at two altitudes using a tethered balloon system for upper ABL sampling and a tower for lower ABL sampling. Sampling was conducted on Byers Peninsula (Livingston Island, South Shetland Islands, Antarctica), a site representative of the marine atmospheric boundary layer (MABL) of the Southern Ocean. Micrometeorological data were recorded concurrently at all sampling heights. In the balloon system, the microbial collector, meteorological and positioning sensors were mounted together on a gondola equipped with a GoPro camera. All collectors included a tail fin to ensure passive alignment of the inlet with the prevailing wind direction. Insets show the study area and the gondola system. Illustration was created with *Illustrae.co*.tool and it is not at scale deliberately to provide a better view of the collection gondola.

**Fig. S2.**
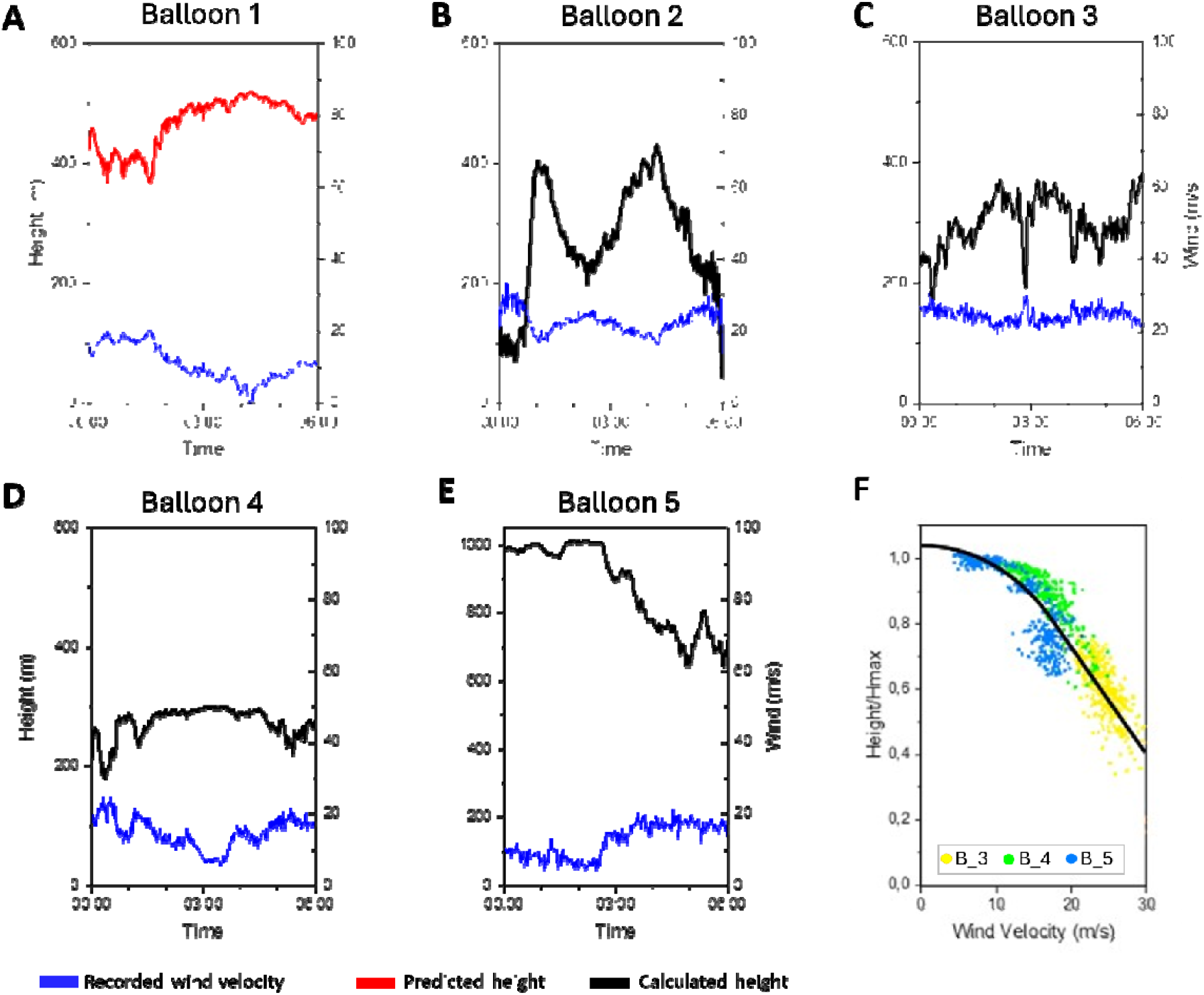
Flight heights and wind velocities of the five balloons. **A-E.** Height reconstructions and wind velocities for balloons 1-5. Black lines indicate balloon heights calculated from atmospheric pressure measurements, whereas blue lines represent the wind speed (WS) recorded at each balloon. Note the different vertical scale in Balloon 5. Because atmospheric pressure data were unavailable for Balloon 1, its height (red line) was predicted using a quadratic relationship between balloon height and wind speed, considering that wind pushed down the tethered balloon. **F.** The relationship between balloon height and wind speed. Continuous line is the fitted quadratic model used to predict the height of balloon 1.

**Fig. S3.**
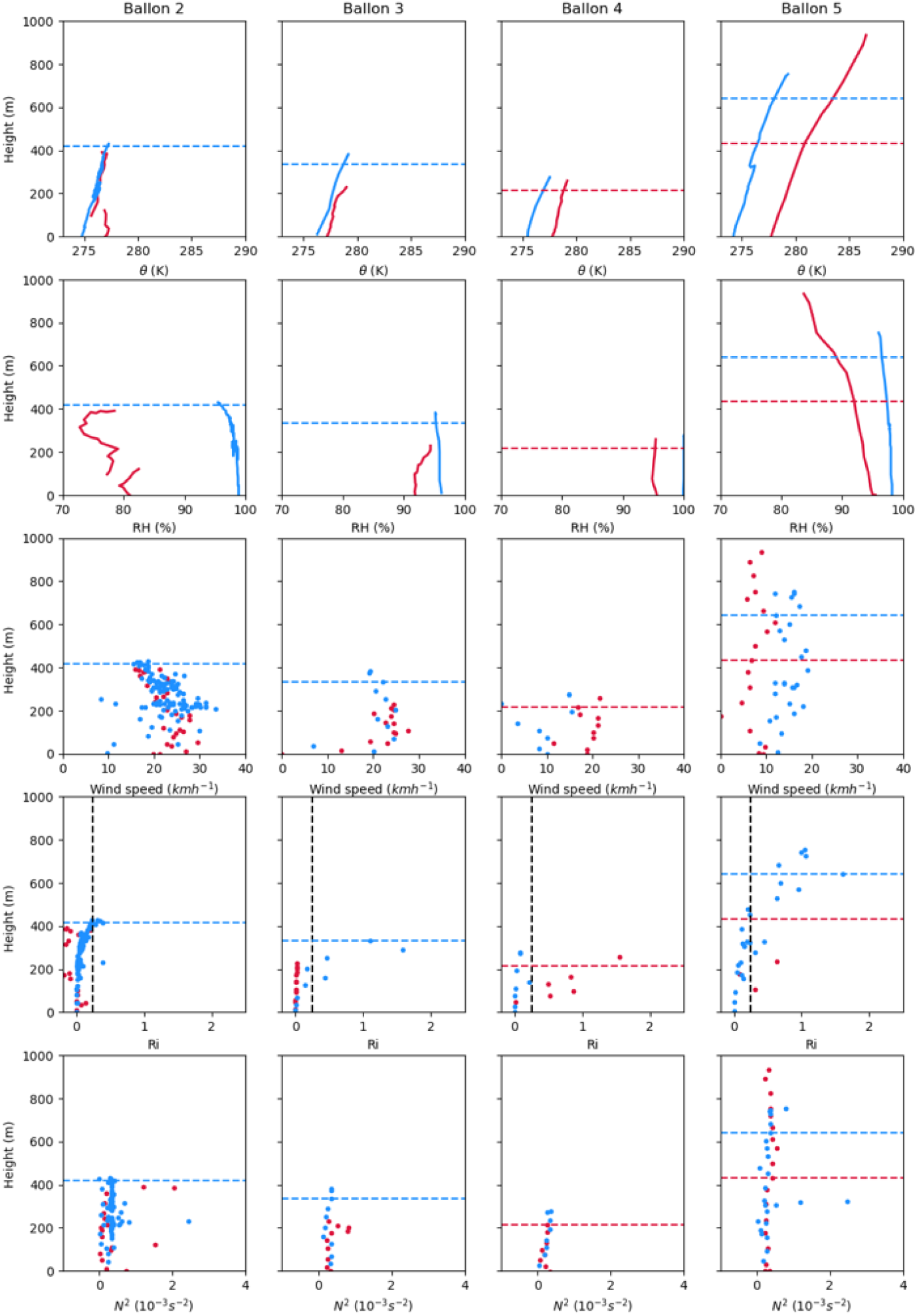
Physical parameters of the ABL as measured from the meteorological instrumentation in the balloons. It includes the potential temperature (θ) in K (first row), relative humidity (RH) in % (second row), wind speed in kmh^−1^ (third row), bulk Richardson number (*Ri*) and stability parameter (N^2^). Plots show the values during the ascending (red) and descending (blue) phase of the profile. The horizontal dashed lines show the position calculated for the ABL heigh . The vertical dashed line indicates the critical *Ri* value of 0.25.

**Figure S4.**
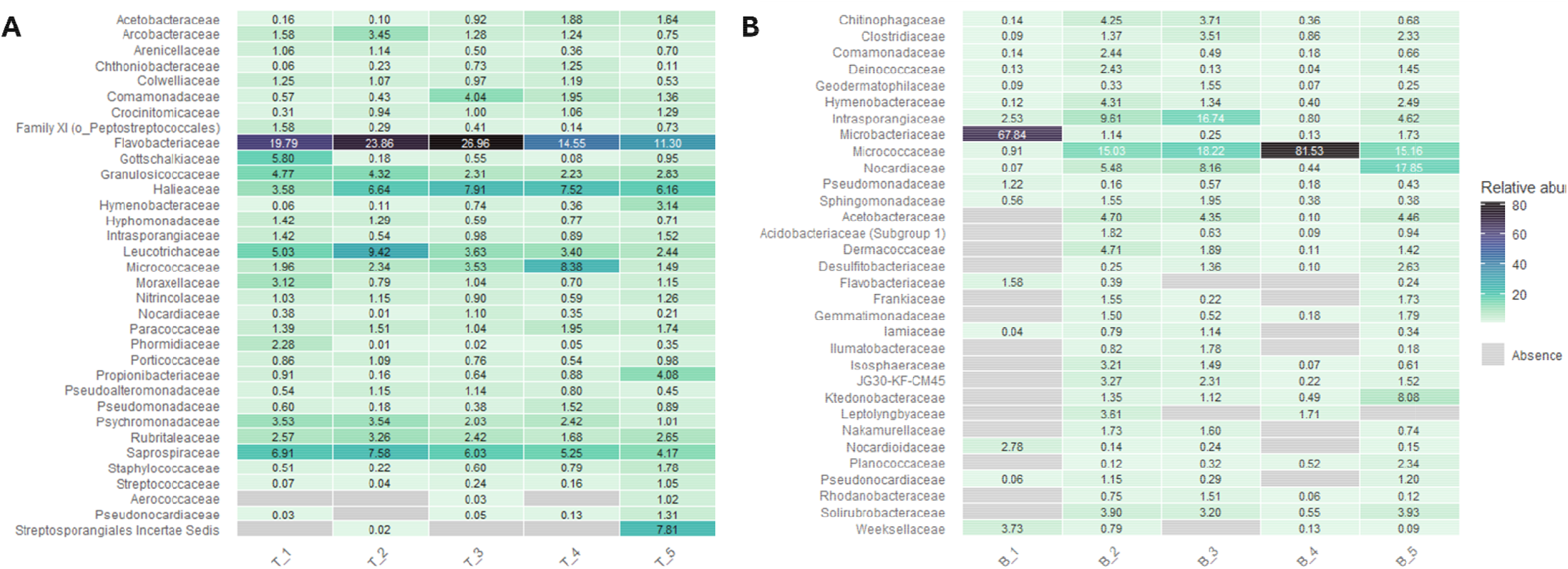
Most abundant families at each ABL layer. **A.** Relative abundance of families (>1% in at least one sample) in lower ABL communities. **B.** Relative abundance of families (>1% in at least one sample) in upper ABL communities. The families are ordered alphabetically.

**Figure S5.**
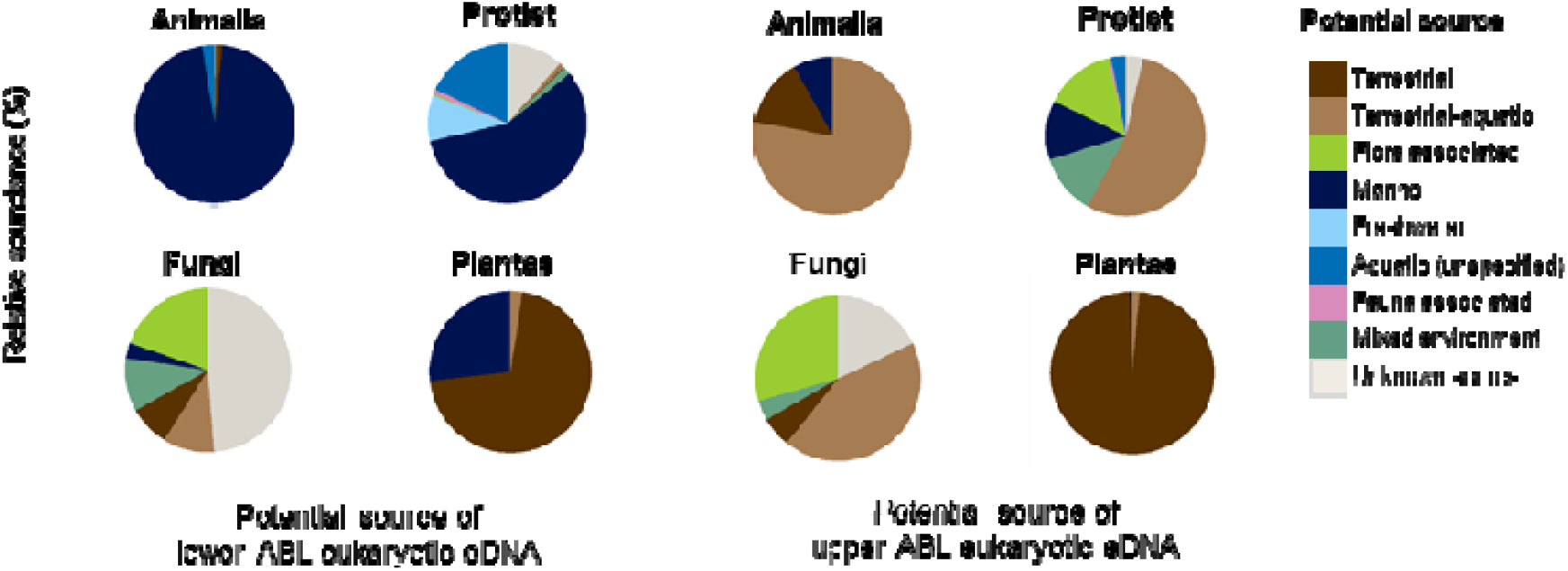
Relative contribution of potential source ecosystems for eukaryotic taxa across ABL layers and kingdoms. Ecosystem assignments were performed at the lowest possible taxonomic level, predominantly at genus level, and at higher taxonomic ranks (e.g., phylum) when finer resolution was not available. *Unknown source* includes taxa that could not be assigned to any ecosystem or that comprise lineages lacking ecological context. See Dataset S7 for detailed source ecosystem annotations.

**Fig. S6.**
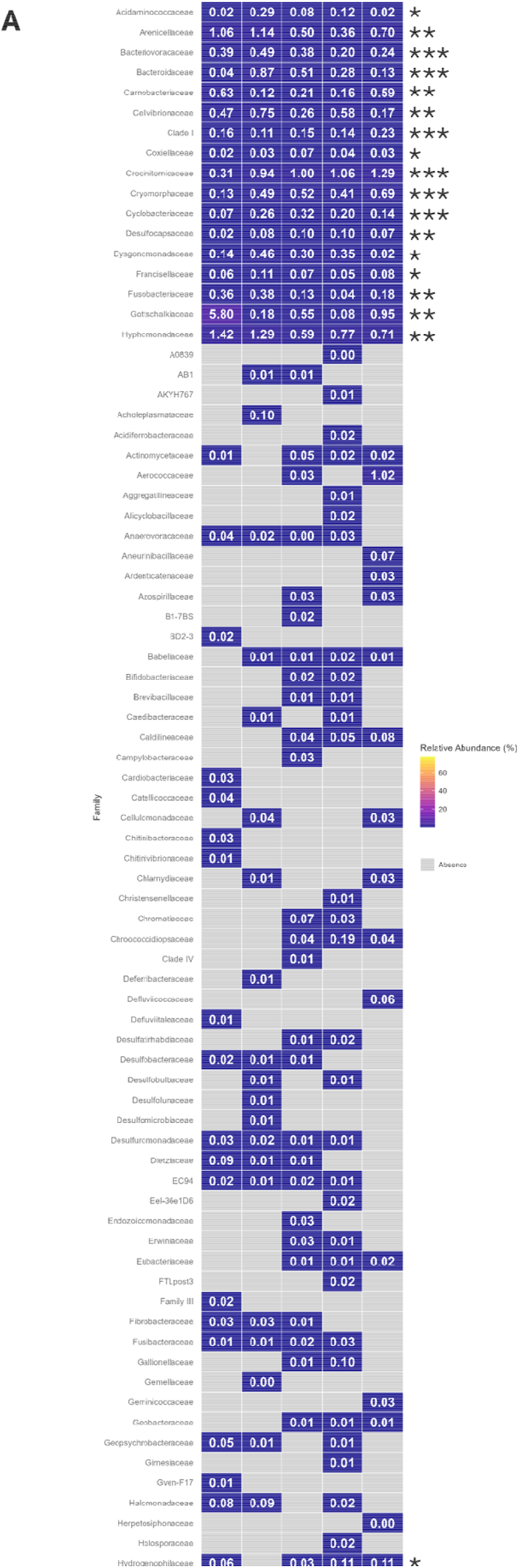
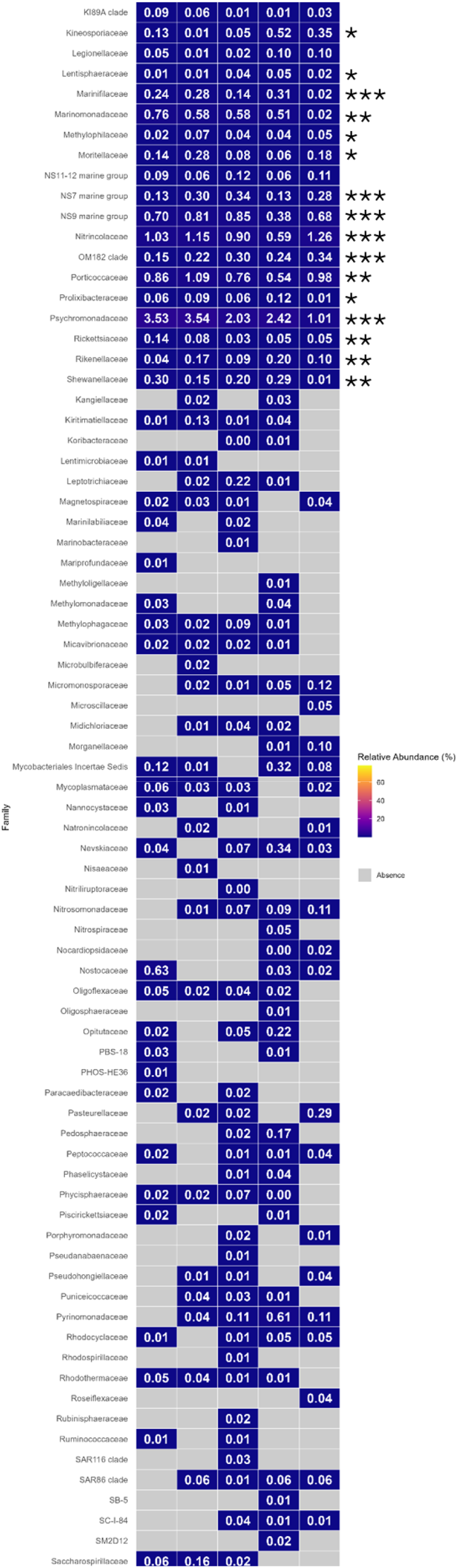
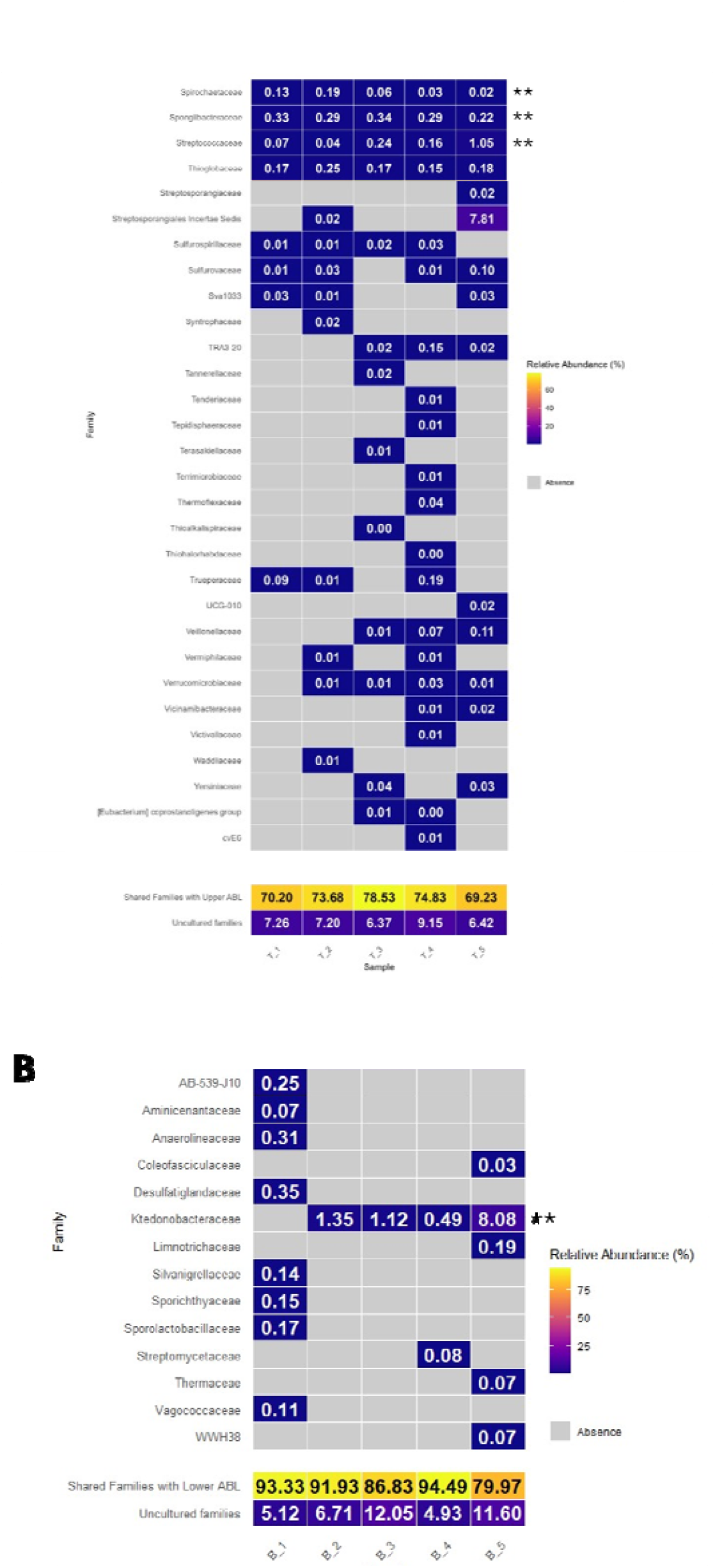
Relative abundance of the bacterial families exclusive to each ABL layer. **A.** Families exclusive to lower ABL samples. **B.** Families exclusive to upper ABL samples. Values of each family in a specific sample displayed inside each tile. The groups *Shared families with upper ABL* (A) and *lower ABL* (B) and *uncultured families* placed at the bottom of each heatmap. **A-B**. Significantly indicator families based on the IndVal test are marked as follows: *** (*p* = 0.001), ** (*p* < 0.01), * (*p* <0.05).

**Fig. S7.**
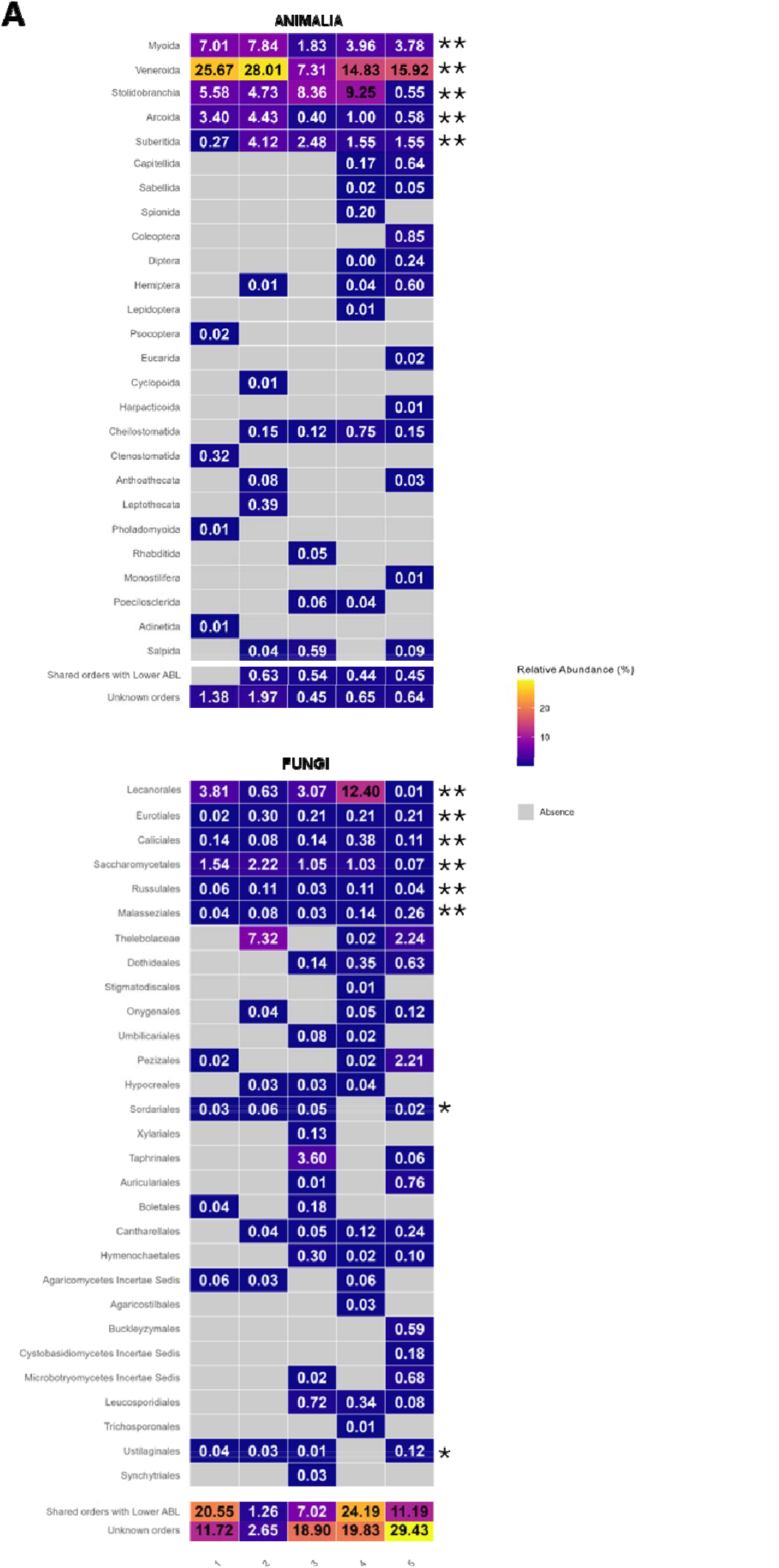
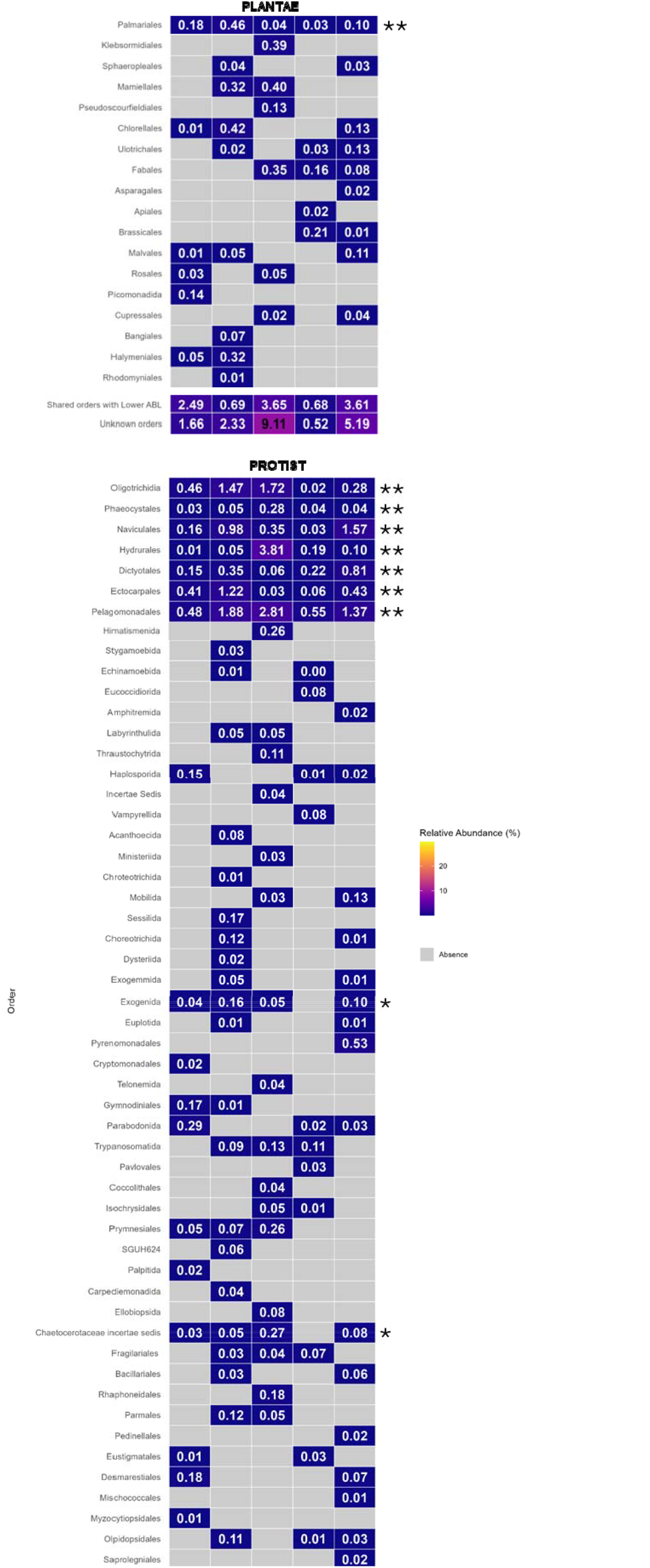
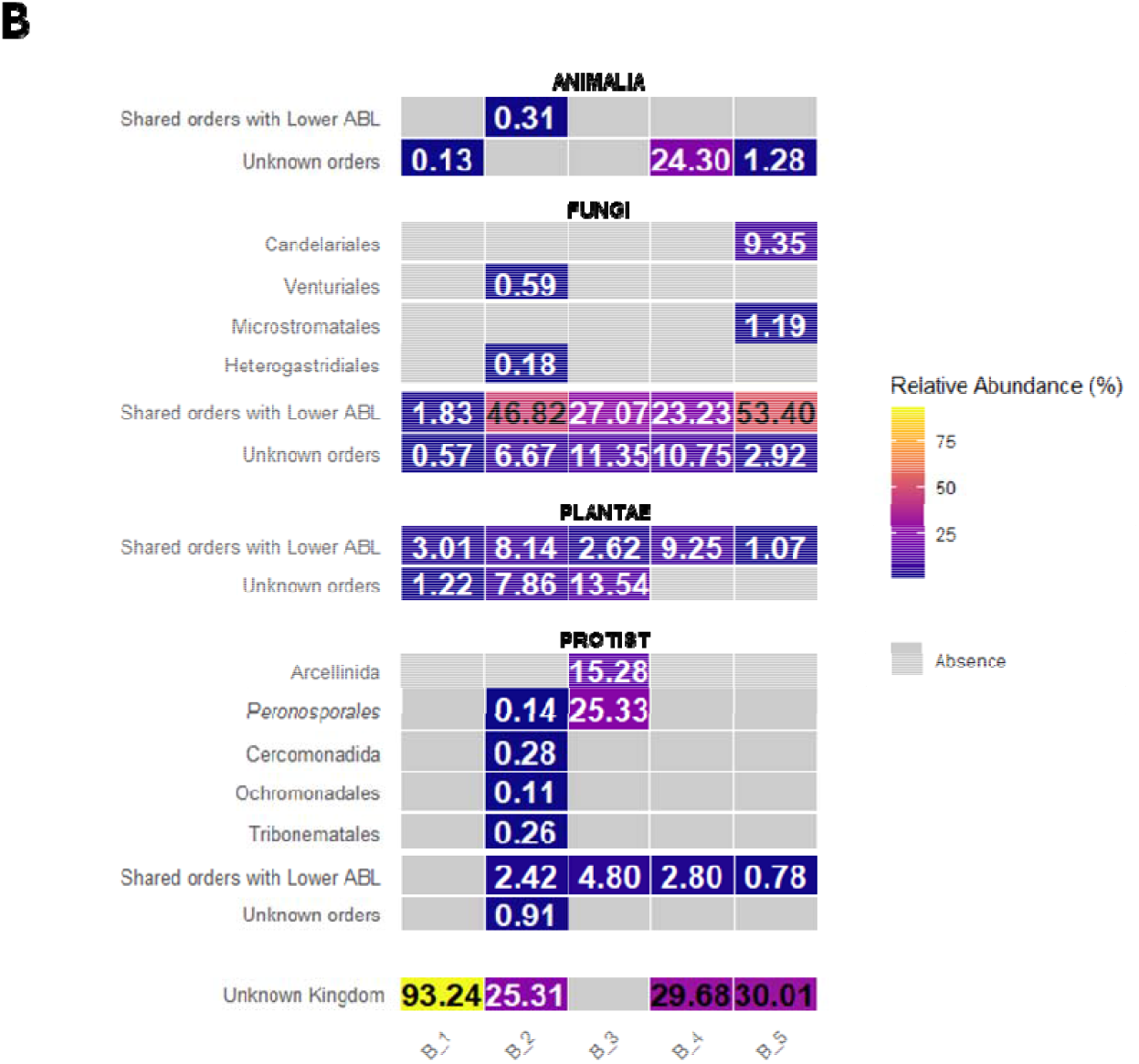
Relative abundance of the eukaryotic orders exclusive to each ABL layer. **A.** Orders exclusive to lower ABL samples. **B.** Orders exclusive to upper ABL samples. Values of each family in a specific sample displayed inside each tile. The groups *Shared orders with upper ABL* (A) and *lower ABL* (B) and *uncultured families* placed at the bottom of each heatmap. **A-B**. Significantly indicator orders based on the IndVal test are marked as follows: ** (*p* < 0.01), * (*p* <0.05).

**Fig. S8.**
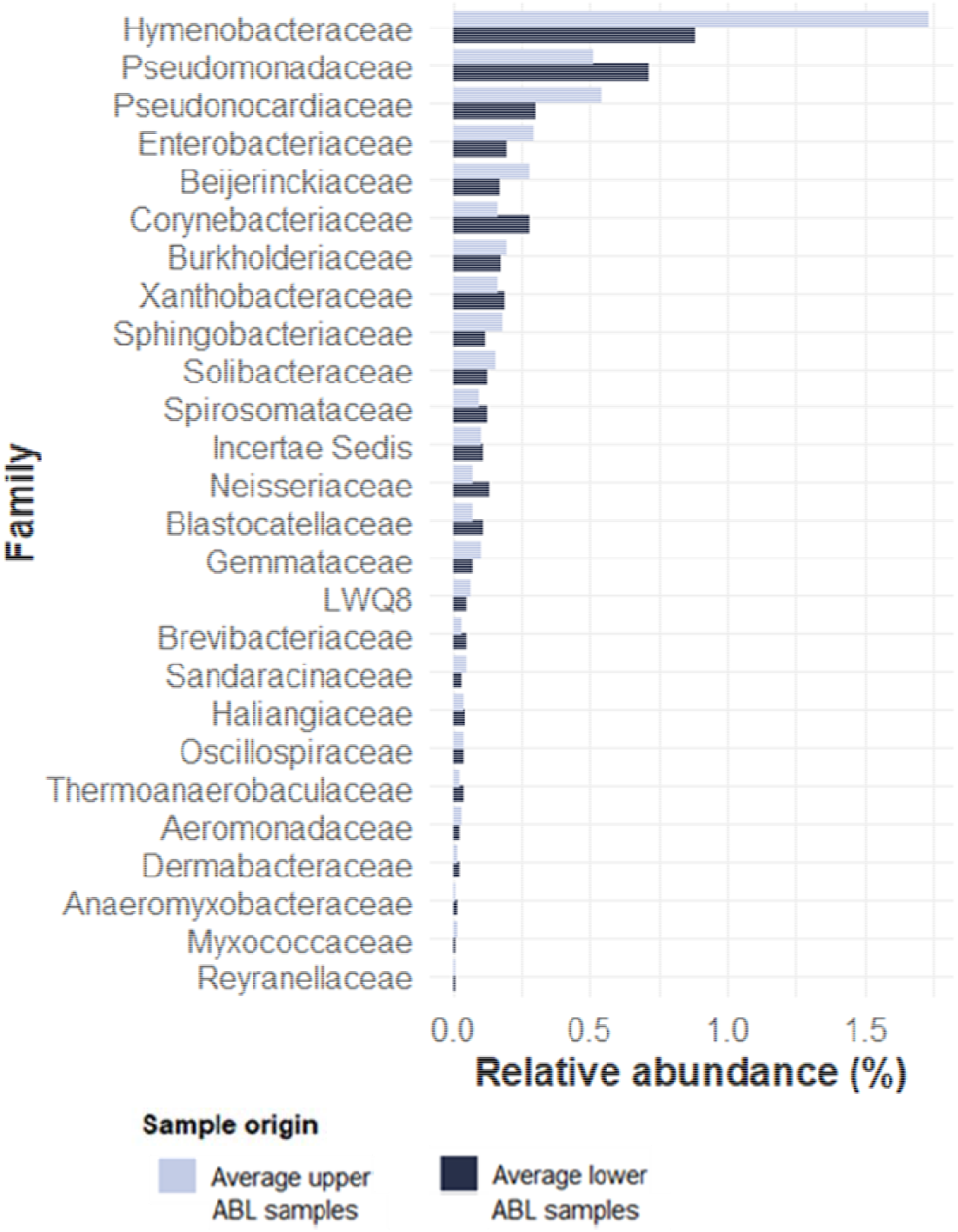
Relative abundance of the bacterial families without clear ABL layer preference. Families were classified as “no ABL layer representative” when the average relative abundance across all sampling events did not exceed twice the average abundance in the opposite layer (neither average upper-layer abundance >2× lower-layer average, nor lower-layer average >2× upper-layer average).

**Fig. S9.**
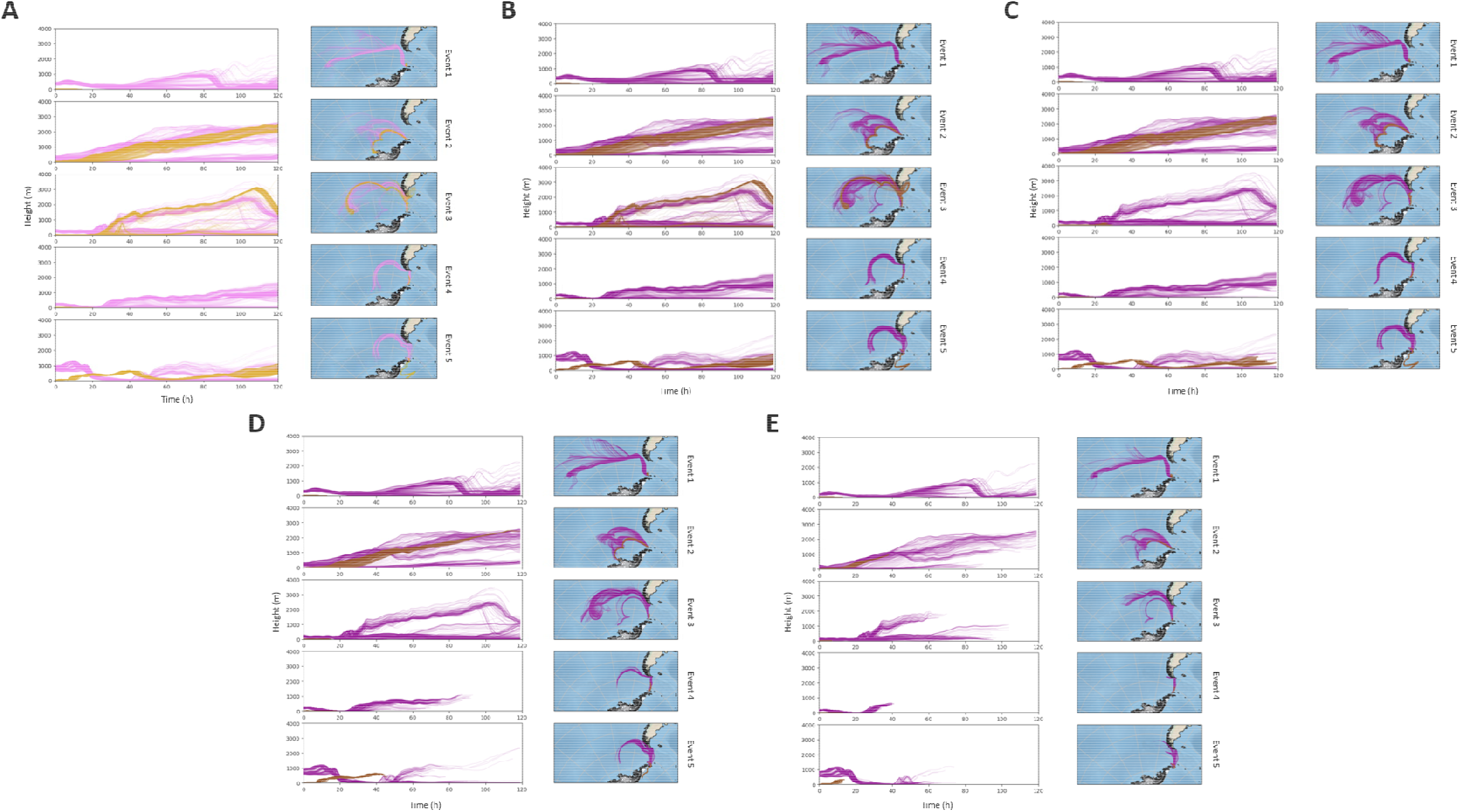
Five-day back trajectories of air masses associated with upper (purple) and lower (orange) ABL samples, showing the temporal evolution of air mass altitude along the trajectories. **A.** Back-trajectories without dry deposition. **B–E.** Back-trajectories with dry deposition considering the ESD of each microorganism: 0.5 μm (**B**), 1.5 μm (**C**), 2.5 μm (**D**), and 4.5 μm (**E**). The particle size classes were considered based on the most abundant microorganisms in the samples. For circular morphotypes, simulations were performed using particles with actual diameters of 0.5, 2.5, and 4.5 μm. For linear morphotypes, three ESD classes were used: 0.5 μm (equivalent to a len. 1.1 μm and diam. 0.3 μm), 1.5 μm (len. 2.5 μm and diam. 1.1 μm), and 2.5 μm (len. 3.4 μm and diam. 1.7 μm). For simplicity, particle sizes are reported as ESD in the figure panels and legend, including those corresponding to circular morphotypes. Simulations were conducted assuming 100% relative humidity and a particle density of 1 g cmO³. When the trajectory height reached to 0 m above the surface, the model is not reliable and the trajectory was interrupted at that point.

**Fig. S10.**
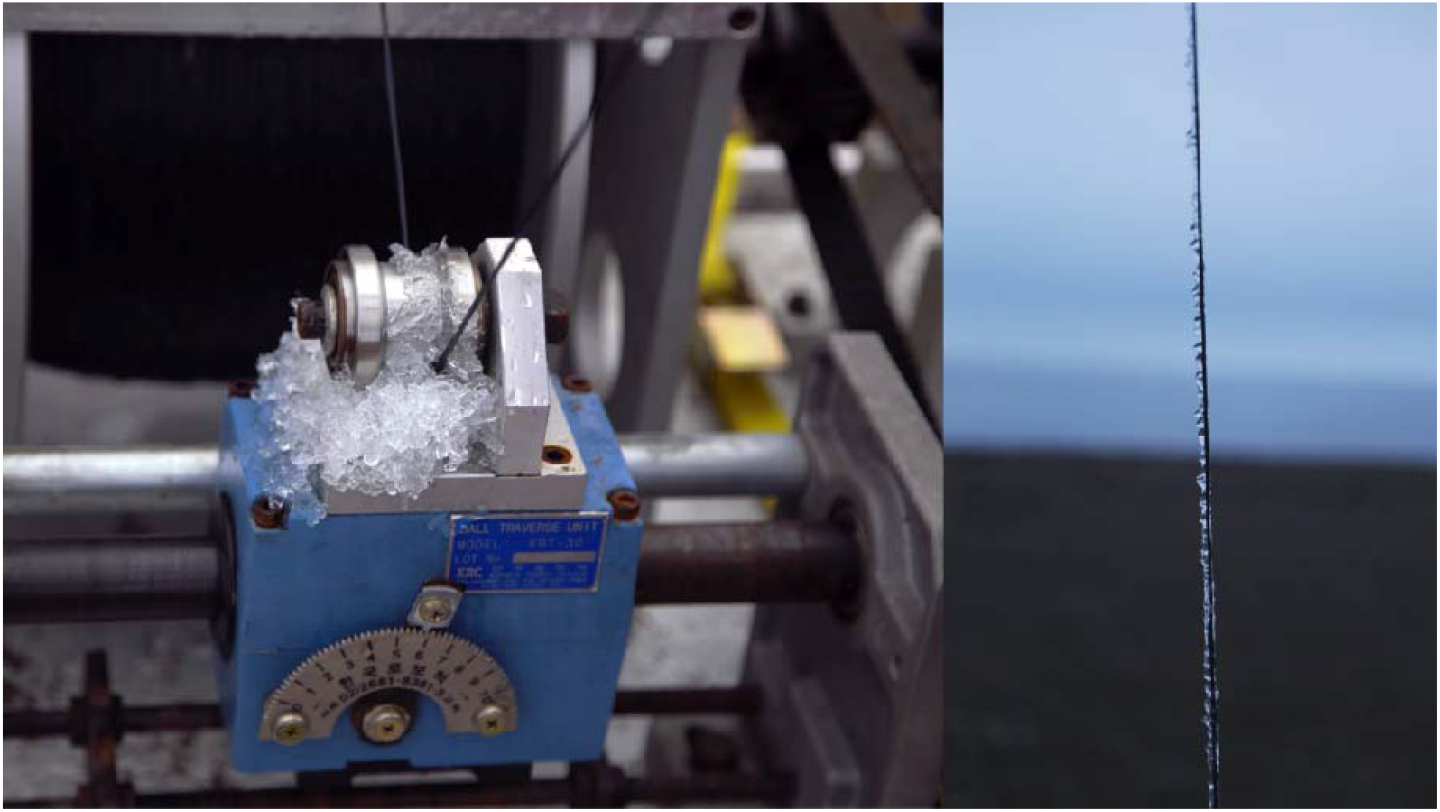
Ice fragments attached to the tether line connecting the balloon gondola holding the collector and the winch. Showing the moment when the balloon descends for sample retrieval during sampling event 5.

**Fig. S11.**
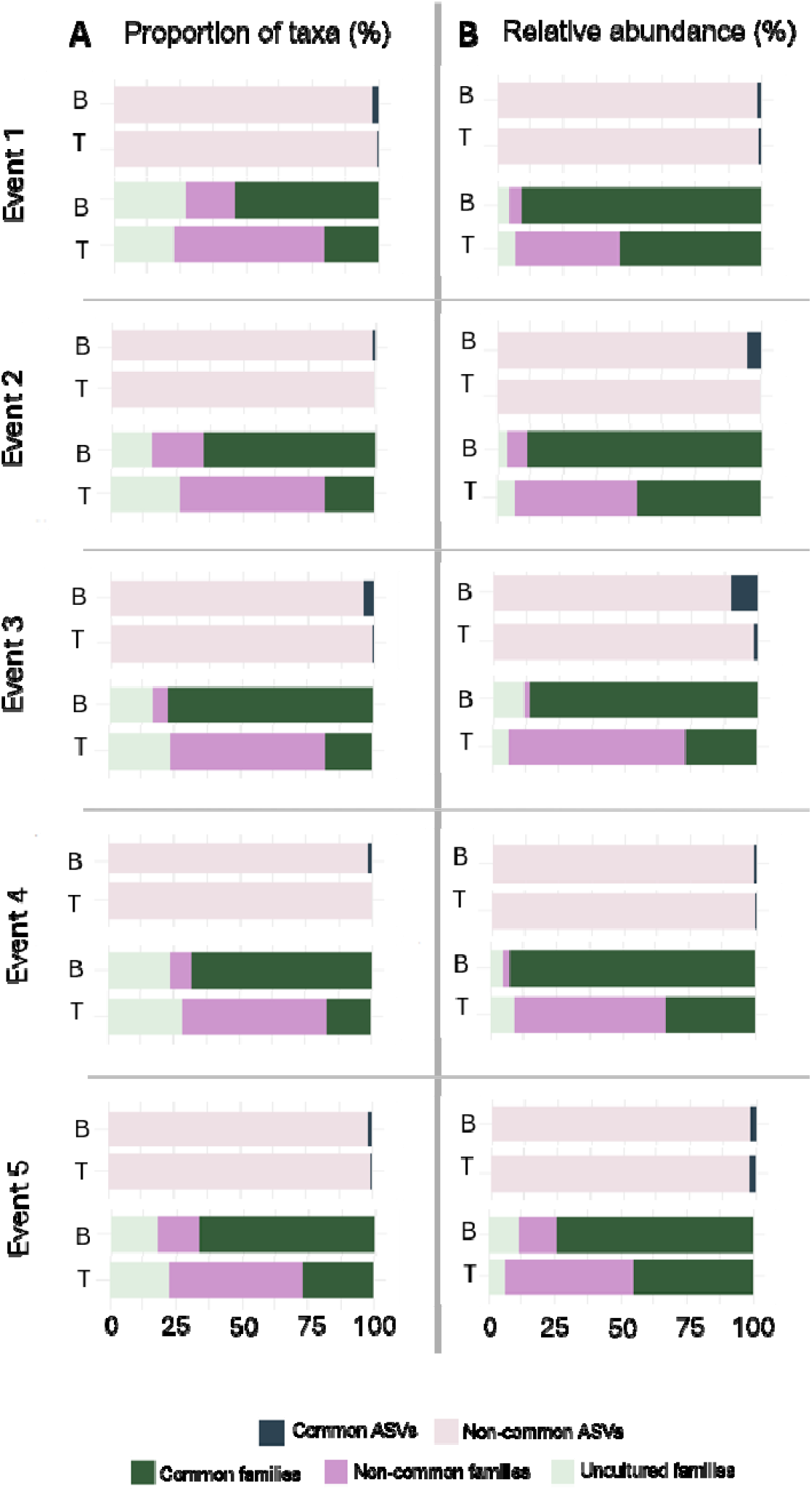
Similarities and differences between upper and lower ABL bacterial communities per sampling event. Proportions of shared and exclusive taxa at the ASV and family levels between upper (B as balloon samples) and lower (T as tower samples) ABL samples are shown for each event. Taxa were defined as shared when detected in both layers. Both proportions of taxa (A) and their relative contributions (B) to community composition are displayed.

## Supplementary Tables

**Table S1.**
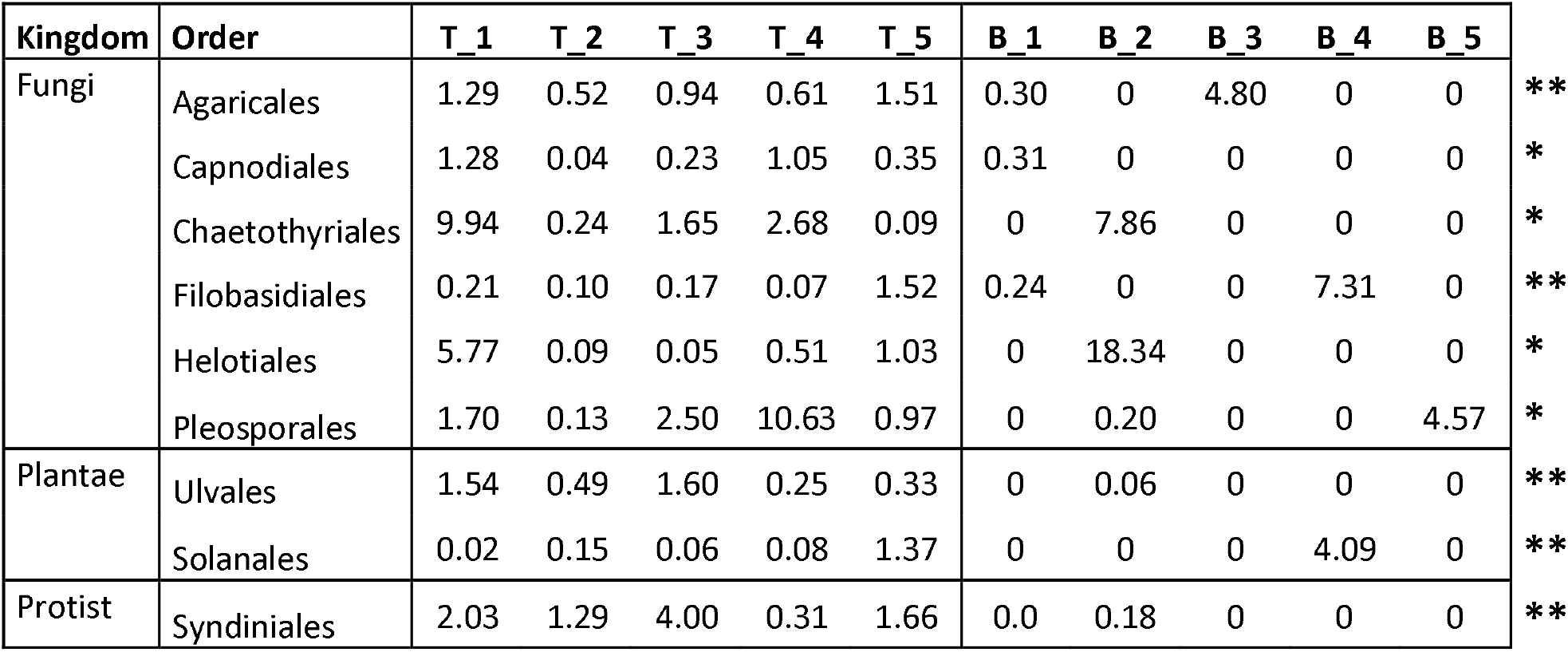
Eukaryotic orders occurring in both lower and upper layers but significantly associated with low-altitude samples according to IndVal analysis. Significant indicator orders are denoted as follows: ** (*p* < 0.01), * (*p* < 0.05).

**Table S2.** ABL height calculated from each balloon during the ascent and descent and sampling region for each balloon. Notice that is not data for balloon 1 to calculate the ABL height.

| Balloon | ABL height (ascent) | ABL height (descent) | Sampling region |
| --- | --- | --- | --- |
| 2 | Not identified (>391 m) | 417 m | 71-430 m (upper ABL) |
| 3 | Not identified (>229 m) | 333 m | 170-382 m (upper ABL) |
| 4 | 215 m | Not identified (>276 m) | 182-299 m (upper ABL) |
| 5 | 433 m | 641 m | 644-1012 m (free atmosphere) |

## Supplementary Dataset Legends

**Dataset S1 - Sample metadata and sequencing accession numbers.** List of all samples, including site locations sampling times, and volumes of air collected, along with the accession numbers for the 16S and 18S rRNA libraries.

**Dataset S2. Balloon micrometeorological recorded measurements and flight height calculated at 1-min intervals for each sampling event.** The dataset includes the meteorological conditions recorded throughout each balloon flight. “Collect_Status” indicates whether the sampling collector was operating or not, allowing the periods of active microbial sample collection to be identified. The forecast flight height (Height_forecast) is represented graphically in Figure S2.

**Dataset S3. Core60 community composition at the ASV level for upper-layer bacterial and eukaryotic assemblages**. For each ASV, the highest confident taxonomic assignment and its relative abundance (%) across sampling events are indicated. The cumulative relative abundance (%) of the Core60 community across sampling events is additionally provided.

**Dataset S4. Potential sources of the most abundant bacterial families in the upper-layer communities**. Families were considered abundant if they reached >1% relative abundance in at least one upper ABL sample (n = 32 families). For each family, the relative abundance of associated genera in each sampling event is shown. Relevant literature consulted (January 2026) is listed to indicate the potential source associated with each genus.

**Dataset S5. Core60 community composition at the ASV level for lower-layer bacterial and eukaryotic assemblages**. For each ASV, the highest confident taxonomic assignment and its relative abundance (%) across sampling events are indicated. The cumulative relative abundance (%) of the Core60 community across sampling events is additionally provided.

**Dataset S6. Potential sources of the most abundant bacterial families in the lower-layer communities**. Families were considered abundant if they reached >1% relative abundance in at least one lower ABL sample (n = 35 families). For each family, the relative abundance of associated genera in each sampling event is shown. Relevant literature consulted (January 2026) is listed to indicate the potential source associated with each genus.

**Dataset S7. Potential sources of the eukaryotic taxa in the lower- and upper-layer communities**. For each taxa, the relative abundance of associated genera in each sampling event is shown. Taxa are grouped by kingdom and ordered alphabetically within each group. Relevant literature consulted (February 2026) is listed to indicate the potential source associated with each taxon. Samples labelled T_1-T_5 correspond to the lower ABL samples, whereas B_1-B_5 correspond to the upper ABL samples.

**Dataset S8. ASVs assigned to the Ktedonobacteraceae family identified in this study were compared with sequences available in the NCBI database.** The search was conducted on 22 November 2025 and only matches with ≥98% sequence similarity are reported. GenBank accession numbers, percent sequence similarity, and geographic metadata are provided. The relative abundances of each ASV in the upper-layer samples are also shown.

## Supplementary Video Legends

**Video S1.** Balloon ascent during sampling event 5, showing the moment when the system passes through a cloud layer. Footage was recorded with a GoPro camera mounted beneath the gondola housing the collector and instrumentation. The collector was not active during the ascent phase.

**Video S2.** Sampling above the cloud layer during sampling event 5. Footage was recorded with a GoPro camera mounted beneath the gondola housing the collector and instrumentation.

---

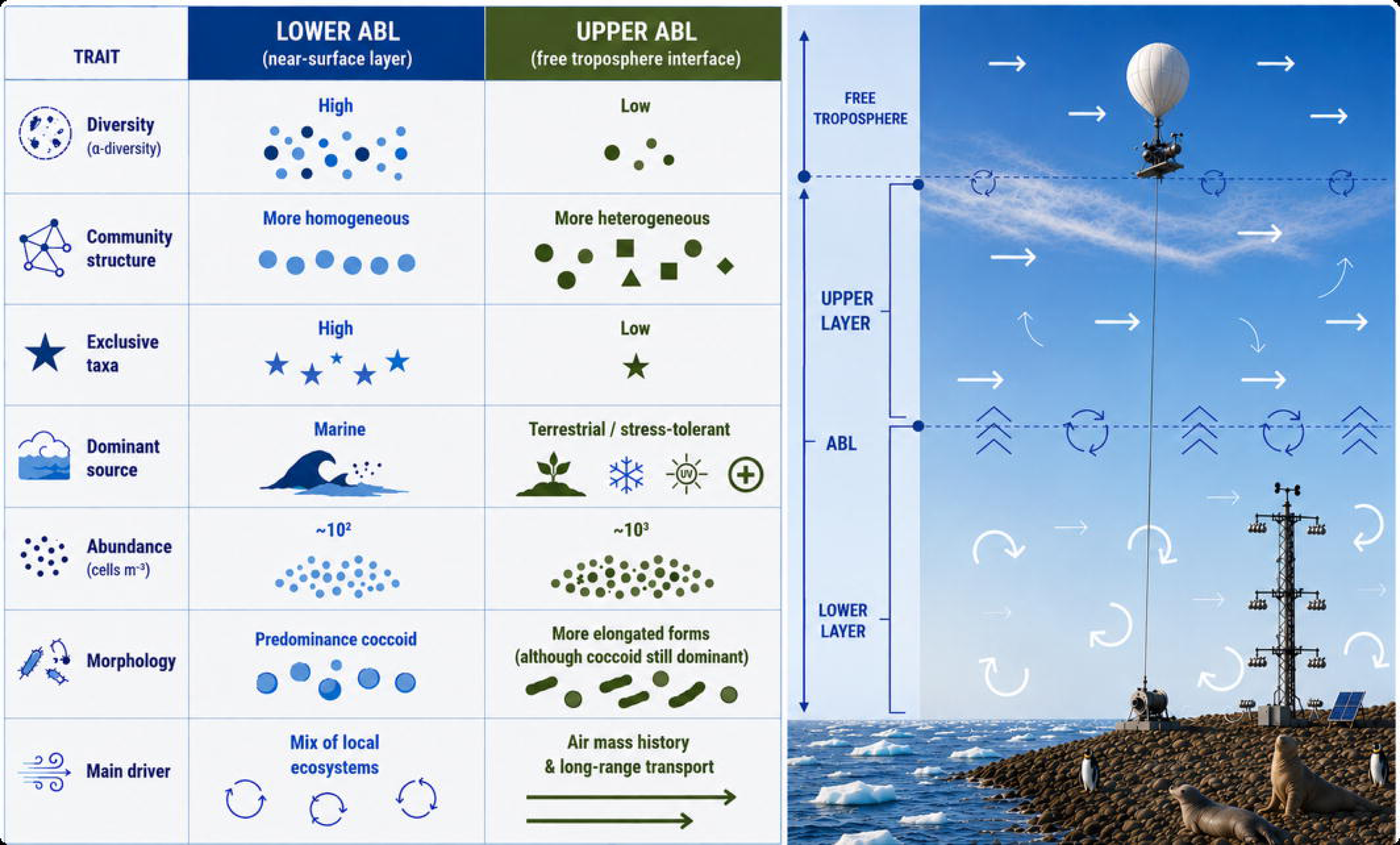

## Notes

### Competing Interest Statement

The authors have declared no competing interest.

